# debar, a sequence-by-sequence denoiser for COI-5P DNA barcode data

**DOI:** 10.1101/2021.01.04.425285

**Authors:** Cameron M. Nugent, Tyler A. Elliott, Sujeevan Ratnasingham, Paul D. N. Hebert, Sarah J. Adamowicz

## Abstract

DNA barcoding and metabarcoding are now widely used to advance species discovery and biodiversity assessments. High-throughput sequencing (HTS) has expanded the volume and scope of these analyses, but elevated error rates introduce noise into sequence records that can inflate estimates of biodiversity. Denoising —the separation of biological signal from instrument (technical) noise—of barcode and metabarcode data currently employs abundance-based methods which do not capitalize on the highly conserved structure of the cytochrome *c* oxidase subunit I (COI) region employed as the animal barcode. This manuscript introduces debar, an R package that utilizes a profile hidden Markov model to denoise indel errors in COI sequences introduced by instrument error. In silico studies demonstrated that debar recognized 95% of artificially introduced indels in COI sequences. When applied to real-world data, debar reduced indel errors in circular consensus sequences obtained with the Sequel platform by 75%, and those generated on the Ion Torrent S5 by 94%. The false correction rate was less than 0.1%, indicating that debar is receptive to the majority of true COI variation in the animal kingdom. In conclusion, the debar package improves DNA barcode and metabarcode workflows by aiding the generation of more accurate sequences aiding the characterization of species diversity.

## Introduction

Motivated by global biodiversity decline, conservation policies and strategies are being implemented to mitigate extinction rates (Driscoll *et al.* 2018; Baynham-Herd *et al.* 2018). Accurate assessments of biodiversity and its change over time are critical to support conservation strategies, to remediate environmental damage, and to manage natural resources, but this information is lacking for most ecosystems (Sogin *et al.* 2006; Hajibabaei *et al.* 2016; Hebert *et al.* 2016; D’Souza & Hebert 2018).

DNA barcoding provides a technological solution to the problem of identifying organisms and characterizing biodiversity (Hebert *et al.* 2003; Hubert & Hanner 2015). Instead of identifying specimens through morphological study, standardized DNA regions—termed DNA barcodes—are used to identify specimens belonging to known species and to recognize new taxa. Reflecting advances in sequencing technology, DNA barcode studies are expanding in scale from analyzing single specimens to characterizing bulk samples, an approach termed metabarcoding, as well as multi-marker and metagenomics approaches (Taberlet *et al.* 2012; Cristescu 2014; Hajibabaei *et al.* 2016; Wilson *et al.* 2019). These advances are providing newly detailed information on species diversity in different geographic regions and habitats (Hajibabaei *et al.* 2012; Hebert *et al.* 2016; Delabye *et al.* 2019; Lopez-Vaamonde *et al.* 2019) while also aiding the identification of invasive species (Brown *et al.* 2016; Xu *et al.* 2017), food web analysis (Wirta *et al.* 2014; Kanuisto *et al.* 2017), and environmental monitoring (Hajibabaei *et al.* 2016; Stat *et al.* 2017; Cordier *et al.* 2019).

Despite the broad adoption of DNA barcoding and metabarcoding, a fundamental problem persists. Efforts to quantify biodiversity from barcode and metabarcode data can be strongly affected by analytical methodology (Clare *et al.* 2016; Braukmann *et al.* 2019). For example, if high-throughput sequence (HTS) data are cleaned suboptimally, the estimated number of taxa may be grossly inflated as variation introduced by sequencing (technical) errors are interpreted as biological variation (Hardge *et al.* 2018).

To reduce the impact of technical errors, sequence reads are often clustered into operational taxonomic units (OTUs) at specific identity thresholds (Elbrecht *et al.* 2018). Several software packages have attempted to increase the accuracy of this OTU method by separating biological signal from technical noise (Rosen *et al.* 2012; Callahan *et al.* 2016; Edgar 2016; Amir *et al.* 2017; Elbrecht *et al.* 2018; Kumar *et al.* 2018; Nearing *et al.* 2018). Many standard denoisers, such as DADA2 (Callahan *et al.* 2016), Deblur (Amir *et al*. 2017), and UNOISE (Edgar 2016), utilize cluster-based approaches, custom error models, or pre-clustering algorithms to account for and correct technical errors. Comparative studies have shown that all three of these methods outperform threshold-based OTU-clustering approaches (Nearing *et al.* 2018). It has also been shown that they produce similar estimates of species richness and relative abundance, but significantly different values for alpha diversity (intra-habitat diversity) and the number of unique exact sequence variants (ESVs) (Nearing *et al.* 2018). When a highly conserved protein-coding region, such as cytochrome *c* oxidase subunit I (COI), is employed as the barcode, structural information can be leveraged to improve denoising. The adoption of this approach can improve the accuracy of alpha-diversity estimates and the quality of identified barcode sequences by ensuring barcodes conform to biological reality. Additionally, rare sequences or important intra-species variants need not be discarded based solely on their abundance and can be retained with higher confidence if they conform to the expected gene structure. This latter benefit will be particularly valuable for work on hyper-diverse communities, (e.g. tropical insects) and for analyses of metabarcode data, where uneven sampling is often the norm and the resolution of intra-species variation is challenging (Elbrecht *et al.* 2018; Nearing *et al.* 2018; Braukmann *et al.* 2019; Zizka *et al.* 2020).

Hidden Markov models (HMMs) are probabilistic representations of sequences that allow unobserved (hidden) states to be inferred through the observation of a series of non-hidden states (Durbin *et al.* 1998; Wilkinson 2019). HMMs have been applied widely in the analysis of biological sequences, in areas such as sequence alignment and annotation (Durbin *et al.* 1998; Eddy 1998). Profile Hidden Markov models (PHMMs) are a variant well suited for the representation of biological sequences with a shared evolutionary origin (Durbin *et al.* 1998; Eddy 1998, 2009). They are probabilistic models that contain position-specific information about the likelihood of potential characters (base pairs or amino acid residues) at the given position in the sequence (emission probabilities) and the likelihood of the observed character given the previously observed character in the sequence (transition probabilities). Once a PHMM is trained on a set of sequences, the Viterbi algorithm can be used to obtain the path of hidden states that align the novel sequences to the PHMM (Durbin *et al.* 1998). The Viterbi path is comprised of hidden match states (indicating the observed character matches to a position in the PHMM) and non-match states: either inserts or deletions. In the context of error correction, hidden non-match states identify the most likely positions at which novel sequences deviate from the PHMM’s statistical profile. In this manner, individual sequences can be queried for evidence of insertion or deletion (indel) errors and adjusted in a statistically informed manner. The conserved protein-coding structure of the most common animal barcode gene, COI, and the wealth of available training sequences (Ratnasingham & Hebert 2007) for this region have allowed PHMMs to be successfully applied in the detection of technical errors in novel barcode sequences (Nugent *et al.* 2020). Correction of technical indel errors in data from protein-coding barcode sequences is an important development as it maximizes the likelihood that both the nucleotide and amino acid sequences correspond to the true biological sequence. Mitigation of indels arising due to technical errors also makes sequence reads from a given specimen more directly comparable, allowing low-frequency point mutations to be eliminated when multiple reads are available for a given biological sequence. Here, we aim to extend the use of PHMMs in COI data processing to allow for the sequence-by-sequence correction (denoising) of technical errors.

This study had four primary goals: (1) design a denoising tool for COI barcode data that utilizes PHMMs to identify and correct insertion and deletion errors resulting from technical error; (2) test the tool’s performance and optimize its default parameters by denoising a set of 10,000 barcode sequences with artificially introduced indel errors; (3) develop, implement, and evaluate a workflow for denoising DNA barcode data produced through single-molecule, real time (SMRT) sequencing of 29,525 specimens on the Sequel platform (Pacific Biosciences); and (4) denoise a DNA metabarcode mock community data set using debar and evaluate the improvement in quality of consensus sequences and the ability to resolve intra-OTU haplotype variation. The denoiser resulting from this work, debar (DEnoising BARcodes), is a free, publicly available package written in R that is available through CRAN (https://CRAN.R-project.org/package=debar) and GitHub (https://github.com/CNuge/debar).

## Materials and Methods

### Implementation

The debar utility includes several customizable steps which denoise DNA barcode and metabarcode data (Figure 1; Supplementary File 1). Corrections with debar are based upon the comparison of input sequences with a nucleotide-based profile hidden Markov model (PHMM) (model training detailed in Nugent *et al.* 2020) using the Viterbi algorithm (Durbin *et al.* 1998). Briefly, debar’s PHMM was trained using a curated set of 11,387 COI-5P barcode sequences obtained from the Barcode of Life Data Systems (BOLD: www.boldsystems.org) public database that were checked to ensure: (i) the sequence was >600 bp in length, (ii) taxonomy was known to a genus level, (iii) there were no missing base pairs, (iv) the amino acid sequence did not contain stop codons, and (v) BOLD’s internal check for contaminants was negative (Nugent et al. 2020). The Viterbi path produced through alignment of the sequence to the PHMMs is used to match the input sequence to the PHMM (by finding the first set of 10 consecutive match states which indicate the absence of indels for the given 10 base pairs). The read is then adjusted to account for detected insertions or deletions (Figure 1). Three consecutive nucleotide insertions or deletions are permitted (not adjusted) as sequences of this kind are more likely to reflect true biological variants than technical errors (they do not result in reading frame shifts and may reflect an insertion or deletion of an amino acid in a functional protein-coding gene). The probability of such changes through sequencing error is relatively low (i.e. for the Pacific Biosciences Sequel platform the baseline probability of three consecutive deletions would be 0.05% (baseline delete probability) cubed, or 0.000125%).

**Figure 1.**
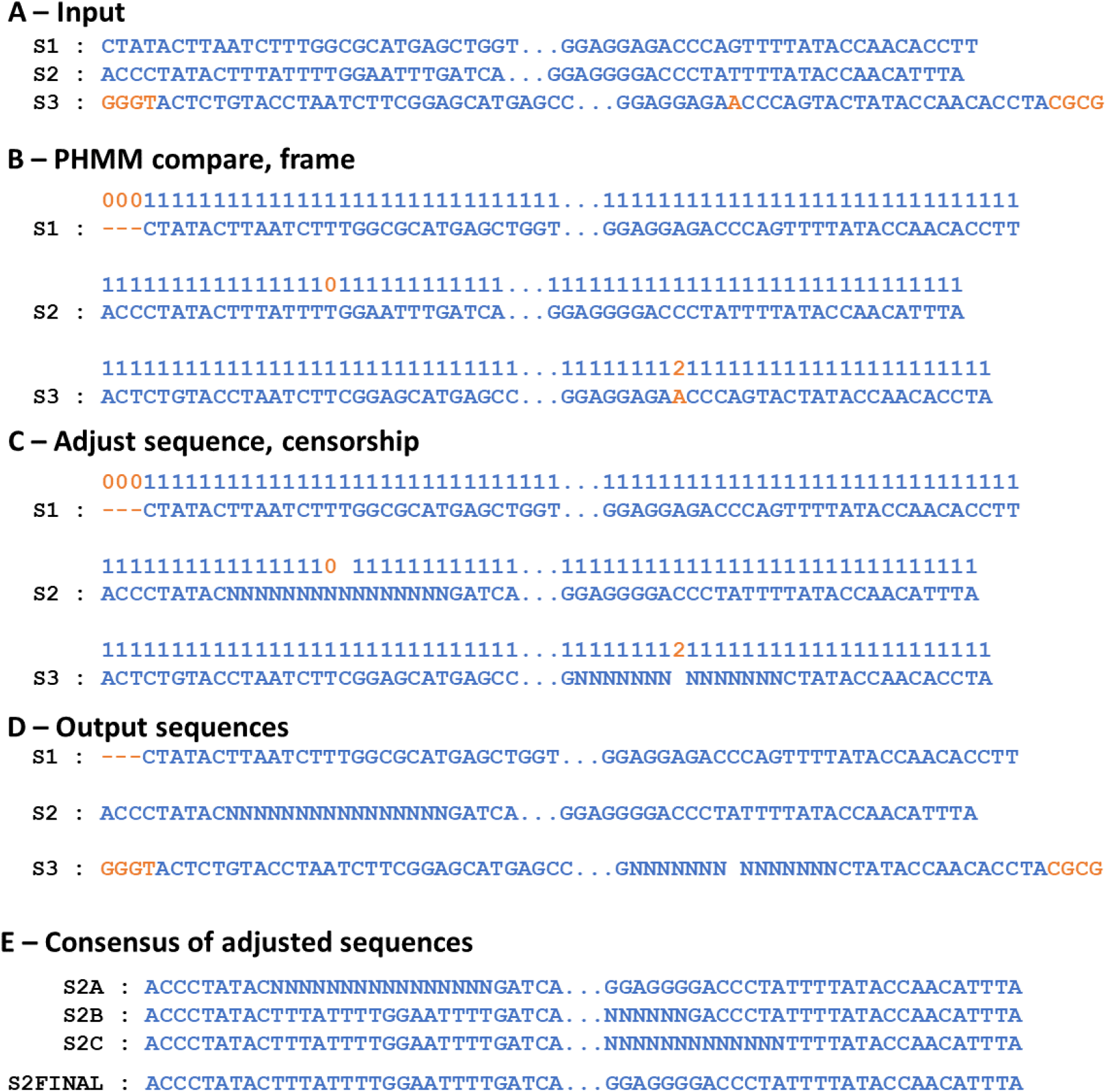
Diagram demonstrating the debar package’s denoising workflow. Blue indicates nucleotides that are part of the barcode region and orange nucleotides in bold font indicate technical errors or sequence from outside of the barcode region. **A.** The debar package operates on a sequence-by-sequence basis, taking each input and constructing a custom DNAseq object. A DNAseq object can receive a DNA sequence, an identifier, and optionally a sequence of corresponding PHRED quality scores. Although not utilized in the denoising, indel-correcting adjustments to the sequence are applied to the PHRED scores as well, so that quality information can be carried from input to output. **B.** Following DNAseq object construction, the sequence is compared to the PHMM using the Viterbi algorithm. By default, the full length (657bp) COI-5P PHMM contained in debar is used to evaluate the sequence. When required, a user may pass a custom PHMM corresponding to a subsection of the COI-5P region (specified using the coil package’s subsetPHMM function) or a custom PHMM trained on user-defined data (Wilkinson 2019). The frame function isolates the correction window, which is the section of the sequence matching the PHMM (the first 10 consecutive base pairs matching to the PHMM on the leading and trailing edges of the sequence establish the section of the input on which subsequent corrections are applied). **C.** The adjust function traverses the section of the sequence and Viterbi path defined by the frame function. When evidence of an inserted base pair (‘2’ label in the Viterbi path) is encountered, the corresponding base pair is removed. When evidence of a deleted base pair is encountered (a ‘0’ label in the Viterbi path) a placeholder ‘N’ nucleotide is inserted. Exceptions are made for triple inserts or triple deletes (three consecutive ‘0’ or ‘2’ labels), which are skipped by the adjustment algorithm, as they are indicative of mutations that would not have a large impact on the structure of the protein-coding gene region and could reflect biological amino acid indels. The total number of adjustments made by debar is limited by the parameter ‘adjust_limit’ (default = 5), sequences requiring adjustments in excess of this number are flagged for rejection, as this high frequency of indels is likely not the result of technical error, but rather other sources of noise such as pseudogenes. Following adjustment, a mask of placeholder ‘N’ nucleotides is applied to base pairs flanking the corrected indel (default is 7bp in each direction, see Figure 3. For derivation of default). Masking of 7bp flanks adjacent to each correction allows imprecise corrections to effectively correct sequence length and also mask true indel locations in the majority of instances. **D.** Following adjustment, the denoised sequences are output by debar. By default, the outputs will include trailing sequence outside of the correction window. Leading information outside of the correction window is dropped, so that sequences are aligned with a common starting position. A user can choose to keep only the correction window, or have both flanking regions appended back on to the sequence output. **E.** If multiple denoised sequences are available (for either a given specimen in the case of barcoding or a given OTU in metabarcoding) then the consensus of the denoised sequences can be taken. The consensus function assumes the sequences have been denoised and their left flanks removed; as a result, they are aligned to one another. The modal base pair for each position is then taken to generate a consensus sequence, and in the case of ties, a placeholder “N” character is added to the consensus.

The denoising of sequences with debar is controlled using a suite of parameters (Figure 1). The censorship parameter is most important as it controls the size of the masks (substitution of nucleotides for placeholder N characters) applied around sequence adjustments. This option is designed to prevent the introduction of errors that would be caused if the denoising process deleted the wrong base pair or inserted a placeholder in the incorrect position. Derivation of the default value for the censorship parameter is detailed in the Methods and Results sections. The package also enables the translation of denoised sequences to amino acids to confirm that denoised outputs conform to the expected properties of the protein-coding gene region. Because debar can interface directly with fasta and fastq files, it enables file-to-file denoising in addition to denoising within an R programming environment. The default PHMM used for denoising by debar represents the complete 657bp barcode region of COI. The package also permits the use of customized PHMMs provided by a user, which allows the denosiser to be applied to data from other gene regions or for the denoiser to be targeted to a specific user-defined subsection of the COI barcode. Training of a PHMM for a new barcode or gene is supported by the R package aphid (Wilkinson 2019), while sub-setting of debar’s default PHMM is enabled by the R package coil (Nugent et al. 2020). Details of the package’s components together with a demonstration of its implementation is available in the package’s vignette (Supplementary File 1).

### Quantification of package performance

#### Simulated error data

The debar package was tested using a phylogenetically stratified random sample of publicly available COI-5P sequences with artificially introduced indels. This test was designed to assess the accuracy of sequence corrections and to obtain a quantitatively informed set of default parameters for the denoising process. A random sample of 10,000 animal COI-5P sequences (excluding those used in PHMM model training) were obtained from BOLD and cleaned using the steps described in Nugent *et al.* 2020 (methods section – BOLD data acquisition). Errors were introduced into each sequence in accordance with the statistical error profile of the Pacific Biosciences Sequel based upon the error profile for COI barcode region in Hebert *et al.* (2018). This profile indicated a baseline indel rate of 0.1% (insertions and deletions equally likely), a baseline substitution rate of 0.5%, and an elevated indel rate for long homopolymers (repeat length of 6,7, and 8+ with indel probabilities of 0.75%, 1.2%, and 3.8%, respectively) (Hebert *et al.* 2018). The location of all errors was recorded so that accuracy of subsequent corrections could be evaluated. Sequences were iteratively processed, and errors were limited to a single insertion or deletion error of one base pair in length (with the error introduction process being repeated for the original sequence when more than one indel occurred), which allowed for the accuracy of corrections to be assessed without the need to consider interaction effects.

The resultant sequences, each with one indel, were then denoised with debar (‘denoise’ function, using the parameter censor_length = 0). The outputs of the denoise function were queried to determine the number and location of indel corrections applied by debar. This information was compared to the recorded ground truth error locations to quantify the following: 1) the frequency with which debar located and exactly corrected indels, 2) the miss distance (number of nucleotide positions) between introduced errors and corrections applied in instances where debar did not correct the indel errors in exactly the correct position, and 3) the frequency at which debar applied an incorrect number of sequence corrections (i.e. 0 correction or 2+ corrections). If one correction was made and the distance between the correction and true indel position was 0, then the correction was considered accurate. Corrections were also considered accurate if all base pairs between the correction location and the true indel position were the same (*i.e.* if base pair 2 in the homopolymer “TTTTT” was an insertion, but the 5^th^ T in the sequence was removed by debar, this is functionally an exact correction as the true sequence is restored). All other corrections at inexact positions were considered inaccurate, and the distance (number of positions) between the correction and true indel location was recorded. The mean and standard deviation of the miss distance were determined and used to select the default censor_length parameter for the debar package, equal to the mean miss distance plus 2 standard deviations (censor_length = ceiling(μ_miss_distance_ + (2 × σ_miss_distance_))). This value was selected as it would be expected to avoid the introduction of an error for > 95% of inexact corrections. Sequences where no corrections or multiple corrections were made had their outputs inspected further to determine if other parts of the denoising pipeline (e.g. the check for stop codons in the translated amino acid sequence or trimming of sequence edges in the framing process) removed the error or led to the complete rejection of the sequence.

#### False correction rate

The performance of debar on sequences with no indel errors was also quantified to determine the frequency and cause of erroneous corrections applied to cleaned, publicly available COI-5P barcode sequences with no known technical errors. A random sample of 10,000 sequences from all the animal COI-5P barcode sequences available on BOLD was obtained (Supplementary File meeting the following criteria was obtained: 1) the barcode was publicly available on the BOLD database, 2) the barcode was > 600bp in length, 3) the barcode did not contain missing characters (“N”) in the Folmer region, 4) the corresponding amino sequence did not contain stop codons, 5) the result of BOLD’s internal check for contaminants was negative, and 6) the sequence was not used in PHMM training and the simulated error dataset. Sequences were processed using debar’s denoise function (censor_length = 0). All sequences that had corrections applied, or that were flagged for rejection, were counted and examined in detail to search for evidence of the proximal cause of the false correction. To search for evidence of taxonomic bias, the taxonomy associated with all falsely corrected sequences were tallied at the order level, and manually examined for evidence of bias.

#### Denoising PacBio Sequel data

We quantified the performance of debar on raw DNA barcode sequence data by interfacing with the existing mBRAVE workflow (http://www.mbrave.net) used to process DNA barcode circular consensus sequences (CCS) obtained with the Sequel platform. A custom analysis pipeline (Supplementary File 3) was constructed to analyze and denoise the final set of CCS barcodes produced by the mBRAVE workflow (one CCS per OTU) (Figure 2). The pipeline was designed to search the final barcodes produced by mBRAVE for evidence of indel errors (by considering the translated amino acid sequence with the R package coil (Nugent *et al.* 2020)), denoise all the associated CCS with detected errors using the debar package, and then regenerate a consensus barcode sequence using the denoised data to produce a final, denoised barcode sequence for each specimen (Figure 2).

**Figure 2.**
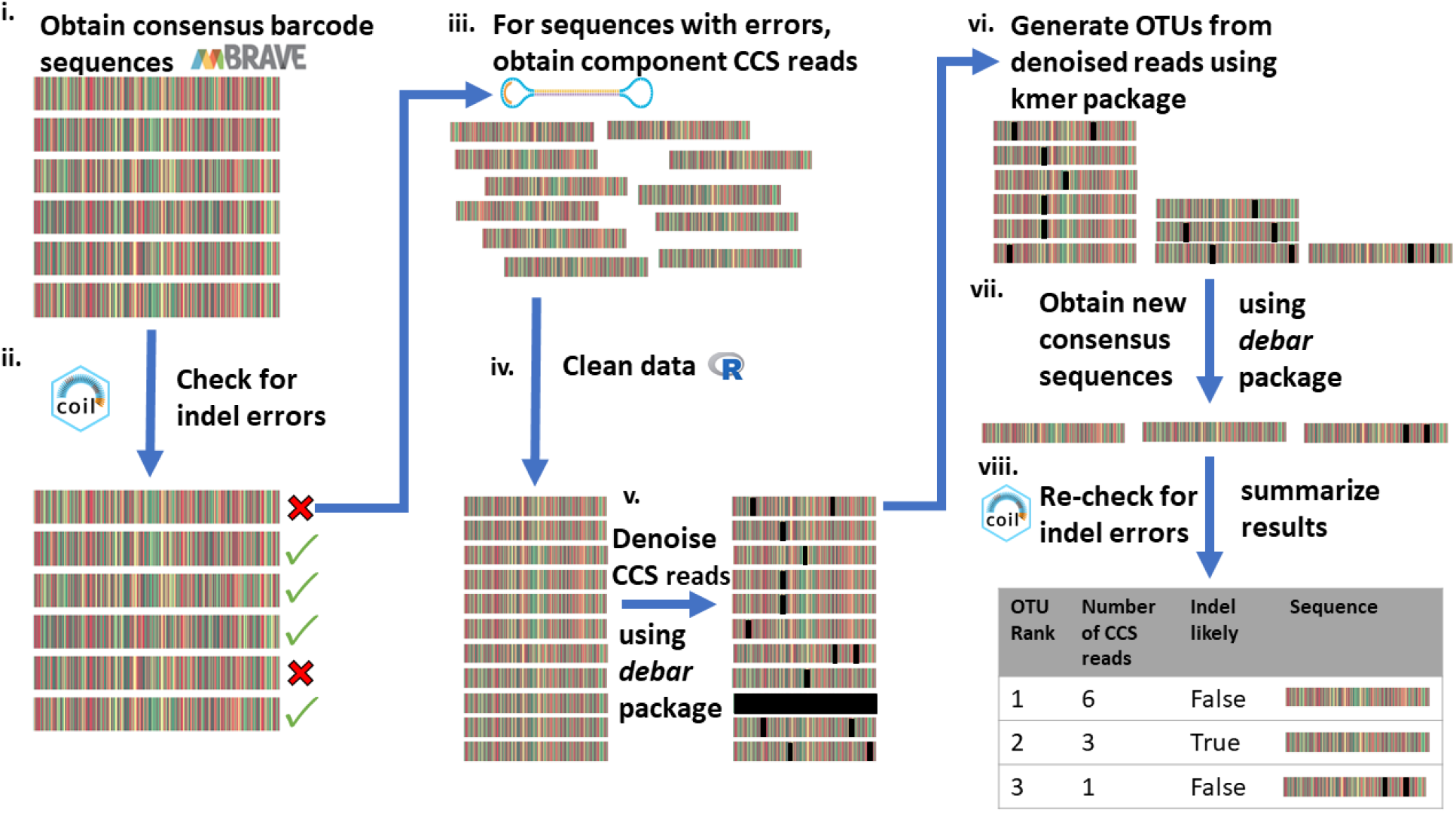
Diagram of the denoising workflow used to improve the quality of barcodes produced by processing Pacific Biosciences Sequel circular consensus data on the mBRAVE platform. **(i)** Pacific Biosciences Sequel data are processed on the mBRAVE platform, and an initial set of barcode sequences is produced. **(ii)** The set of consensus barcode sequences produced by the mBRAVE platform are obtained and analyzed with the coil package, using the ‘coi5p_pipe’ function (default parameters). Sequences displaying evidence of an indel (either the presence of a stop codon when translated to amino acids or an amino acid sequence with a low likelihood score) are retained for further denoising. **(iii)** For each barcode with evidence of an error, all component CCS reads (and associated metadata) derived from the given specimen are obtained from mBRAVE. **(iv)** Based on the mBRAVE metadata, sequences are trimmed to remove primers, MID tags, and adapter sequence. The reverse complement of reads are taken when required. **(v)** The ‘denoise_list’ function of debar is used to denoise all CCS reads (options: dir_check = FALSE, keep_flanks = ‘right’, censor_length = 7). Rejected reads (those flagged by the denoise_list function) are removed from the dataset. **(vi)** For each specimen, the reads are clustered into OTUs using the R package kmer (clustering threshold = 0.975). This is done to mitigate the influence of outlier CCS or contaminant sequences. **(vii)** For each OTU, a consensus sequence is generated using debar’s ‘consensus’ function. For each specimen, OTUs are ranked based on the number of component CCS reads they contain. **(vii)** The consensus sequences are reassessed with coil. If the top-ranked consensus sequence now passes the coil check, it is deemed to have been successfully denoised, and it is selected as the output barcode. If not, the check is repeated for the second-ranked consensus sequence (when available), and this output is retained if it is barcode compliant. If neither the first nor second highest ranked consensus sequence passes the coil check, then the original (pre-denoising process) barcode is retained, as no meaningful improvement was made. In this situation the barcode is flagged as likely to contain an error.

The outputs of this analysis were examined to determine if the debar pipeline decreased the number of technical errors in the barcode sequences and that those barcode sequences resulted in likely amino acid sequences when translated. Initial quantification of the improvement was conducted by comparing the number of barcode sequences whose amino acid sequences were flagged by the R package coil (Nugent *et al.* 2020, default parameters) before and after denoising. Barcodes are flagged by coil when they possess a stop codon when translated to amino acids or when the resultant amino acid sequence is improbable, both indicating that the sequence likely possesses an indel error.

Since the coil and debar packages both employ the same nucleotide profile hidden Markov model (coil also utilizes an amino acid PHMM), an independent test of pipeline effectiveness was also conducted. The effectiveness of the denoising pipeline was quantified by submitting both the original and denoised barcode sequences to BOLD. It was used to determine the number of original barcodes and denoised barcodes with evidence of stop codons after aligning the sequences using the BOLD’s hidden Markov model (a model developed independently of the debar PHMM) and translating the sequence using the appropriate translation table corresponding to the taxonomic information accompanying the sequence record. Comparison of these numbers made it possible to quantify the increase in barcode-compliant sequences (i.e. those with no stop codon) produced by debar. Additionally, the Sequence Quality Report on BOLD was examined to determine the number of unknown nucleotides (“N”) in the barcode sequences after denoising. The report categorizes barcode quality as: high (<1% Ns), medium (<2% Ns), low (<4% Ns), or unreliable (>4% Ns), and the number of barcodes in these different categories was recorded.

#### Denoising metabarcode data

To characterize debar’s performance on metabarcode data, we analyzed a metabarcode dataset for a mock arthropod community (Braukmann *et al.* 2019). These data derived from a single sequencing run on an Ion Torrent S5 on COI amplicons generated by pooled DNA extracts from abdomens from single specimens of 369 arthropod species (methods described in detail in Braukmann *et al.* 2019). Sequences were from a 407bp fragment of the COI barcode region targeted using the primers MlepF1 and LepR1 (Hebert *et al.* 2004; Braukmann *et al.* 2019). Following amplification and sequencing on the Ion S5, quality control, sequence dereplication, chimeric read filtering, matching to reference sequences, and clustering were performed on mBRAVE (Braukman *et al.* 2019). Two sets of data resulted from this process, a set of 123,926 unique sequences that were assigned to 398 different Barcode Index Numbers (BINs) (Ratnasingham and Hebert 2013) through the comparison to reference sequences (matched at >98% similarity), and a set of 2,199 unique sequences not matching to available references that were clustered into an additional 1,255 OTUs at a 97% similarity threshold (using clustering algorithm described in Braukmann *et al.* 2019).

All sequences were denoised using debar’s denoise_list function and a custom nucleotide PHMM. The custom PHMM was a 398bp subset of the complete COI PHMM (PHMM profile positions 250 – 648), corresponding to a segment of the Folmer (Folmer *et al.* 1994) region targeted by the metabarcoding primers. The PHMM was created using coil’s ‘subsetPHMM’ function (Nugent *et al.* 2020). After denoising, two tests were conducted to determine if denoising improved the quality of the metabarcode pipeline’s output data.

First, for each BIN and OTU consensus sequences were generated using denoised sequences and the debar function ‘consensus_sequence’. These consensus sequences were assessed for evidence of stop codons using coil and the same custom PHMMs used in denoising (function coi5p_pipe with the additional parameter: trans_table = 5). This test revealed the number of denoised consensus sequences which contained a stop codon when translated to amino acids, indicating an indel error persisted in the nucleotide sequence. The centroid sequences for the BINs and OTUs were used as a baseline metric for the number of barcode-compliant sequences. For each BIN, centroid sequences were obtained by clustering the sequences in the group using the R package kmer’s ‘otu’ function (parameters: k = 4, threshold = 0.95) (Wilkinson 2018, Version 1.0.0). For the OTUs, centroids were obtained from data generated by mBRAVE. All centroids were assessed with coil (Nugent *et al.* 2020, Version 1.0), and the number of barcode-compliant representative sequences for the original centroids and the final consensus sequences was compared.

Secondly, the individual sequences within each BIN and OTU were analyzed with coil to determine the number that were likely error free, as evidenced by the absence of stop codons after translation. This assessment was repeated on the denoised reads to determine the effectiveness of debar in correcting errors in individual sequences and to reveal if the denoising process improved the resolution of ESVs for subsequent analysis of intra-species genetic variation by placing the ESVs in reading frame and reducing the frequency of identified indel errors.

## Results

### Quantification of package performance

#### Simulated error data

Debar was used to correct 10,000 barcodes, each with a single indel error (Supplementary File 1). The denoised sequences and associated data were compared to the ground truth error locations to determine the accuracy of corrections applied by debar (Figure 3). For 9,459 sequences (95.59%), a single correction was applied by debar, indicating that the package correctly identified the type of error in these sequences. However, debar either failed to recognize an indel or made too many corrections (2+) in the other 541 sequences. No correction was made for most (426) of these sequences, meaning that debar’s PHMM did not identify the indel error. The overlooked indels were largely restricted to the terminal regions of the sequence; 75% (329/426) of them were positioned within 20 base pairs of the read termini (Figure 4), regions that only comprised 5% (40bp/650bp) of the sequences. The cause of this is that the debar denoising algorithm uses the first observation of 10 consecutive bp matching to the PHMM to establish the corrective window. Errors on the periphery of sequences therefore lead to trimming of the sequence (via the keep_flanks function) instead of indel correction. A substantial fraction of the remaining uncorrected indel errors (43) occurred between positions 452 to 465 (Figure 4), a region associated with a 3bp indel present in some animal groups and absent in others. Its presence reduced the PHMM’s indel detection ability in this region due to greater true variability. Not all unidentified indels were retained in the final output sequences as double checks of debar (employing the keep_flanks and aa_check parameters) identified many (266/426 – 62%) of the uncorrected sequences and either omit the problem region or flag the sequence as likely to contain an error. Therefore, debar’s double checks allow many false negatives to be trimmed or flagged as problematic.

**Figure 3.**
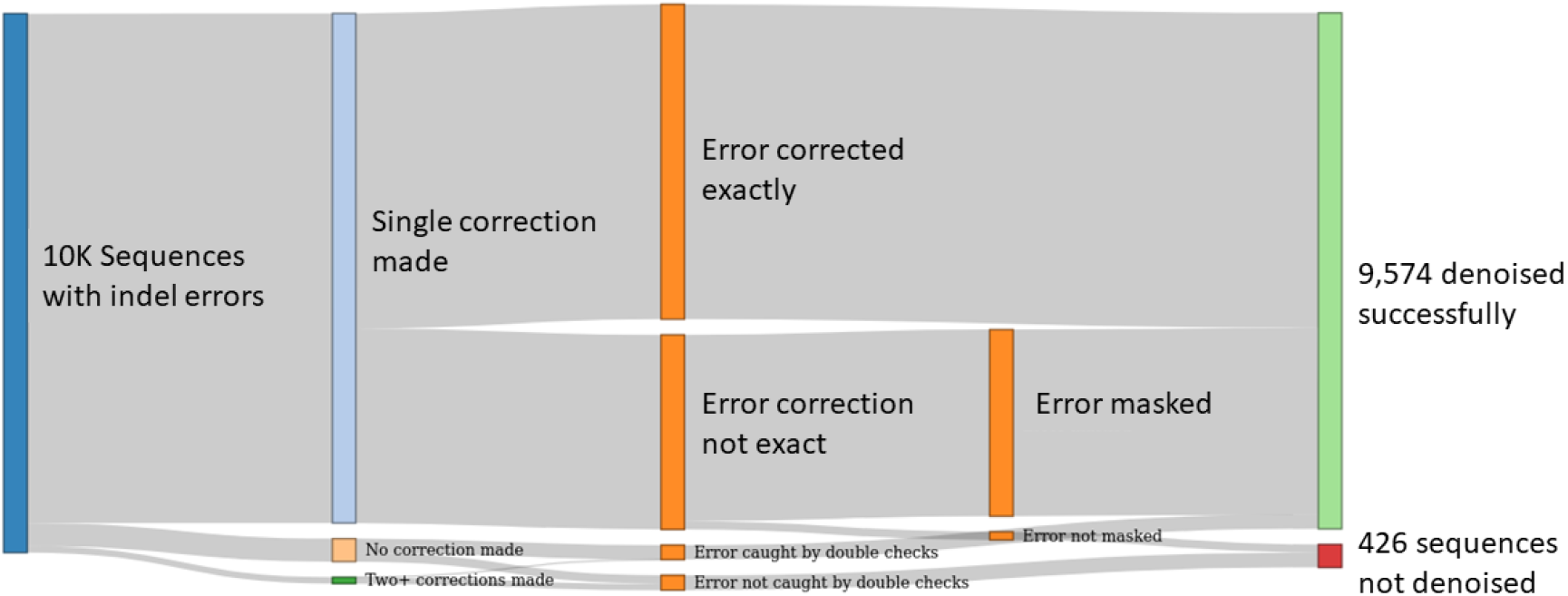
The debar package’s denoising of 10,000 COI sequences containing single insertion or deletion errors. So that exact error positions were known, errors were artificially introduced in accordance with known probabilities for COI DNA barcode data from the PacBio Sequel platform (Hebert *et al.* 2018). Denoising was accomplished through altering sequences in accordance with the Viterbi path yielded by comparison to the PHMM. The correct number of adjustments was made for 9,455 sequences, and 61.8% of these corrections located the indel exactly. Masking of 7bp flanks adjacent to each correction allowed imprecise corrections to correct sequence length and mask the true indel location 96% of the time. For the 545 instances where an incorrect number of adjustments were made, 269 were caught through query of the amino acid sequence for stop codons and the trimming of spurious matches at the edge of sequences. Overall, 95.74% of errors were effectively corrected or identified as erroneous.

**Figure 4.**
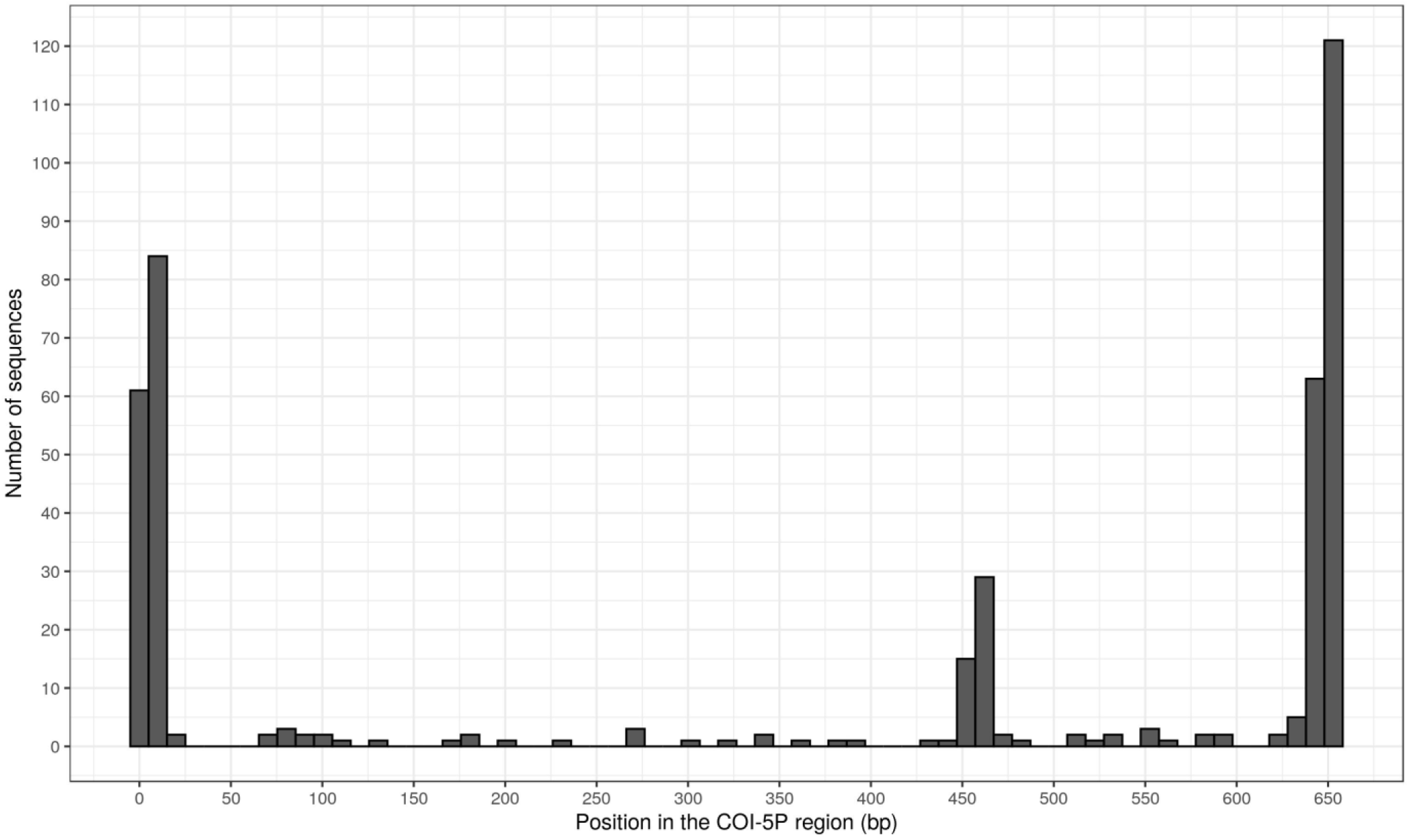
Histogram indicating the position in the COI-5P region of the 426 uncorrected indel errors from the 10,000-sequence artificial error dataset. The x axis indicates the base pair position in the COI-5P profile, and the y axis displays the number of sequences that contained an uncorrected error at the given range of positions (bins of 10 base pair positions).

For 119 sequences (1.2%), two or more corrections were applied by debar when only a single indel existed (Figure 3). In contrast to the false negatives, debar’s double checks only captured three of the false positives. Many of the false corrections appeared to be the presence of indels near codons that are not present in all animals. Due to true biological variation in the training data, these regions of the PHMM have higher probabilities of transitioning from a match state to an insert or delete state, and therefore indels in these locations are sometimes handled incorrectly (i.e. the sequence is characterized as having two deleted base pairs, when there was a 1bp insertion). Because false corrections of this type result in sequences that conform to the structure of the protein-coding gene region (*i.e.* a lack of stop codons in the amino acid sequence), they are not identified by debar’s aa_check function.

The 9,459 sequences for which the presence of a single indel was correctly identified were further analyzed to determine how accurately they were located (Figure 3). The analysis showed that debar was able to exactly locate and correct 5,847 (61.81% of sequences in single correction category) of the indel errors in the dataset. For the other 3,612 sequences (38.19% of the single corrections category), the indel corrections were not placed in exactly the correct position (Figure 5). For these sequences, the average distance between the true indel location and the applied correction was 2.31 base pairs (standard deviation = 1.9767).

**Figure 5.**
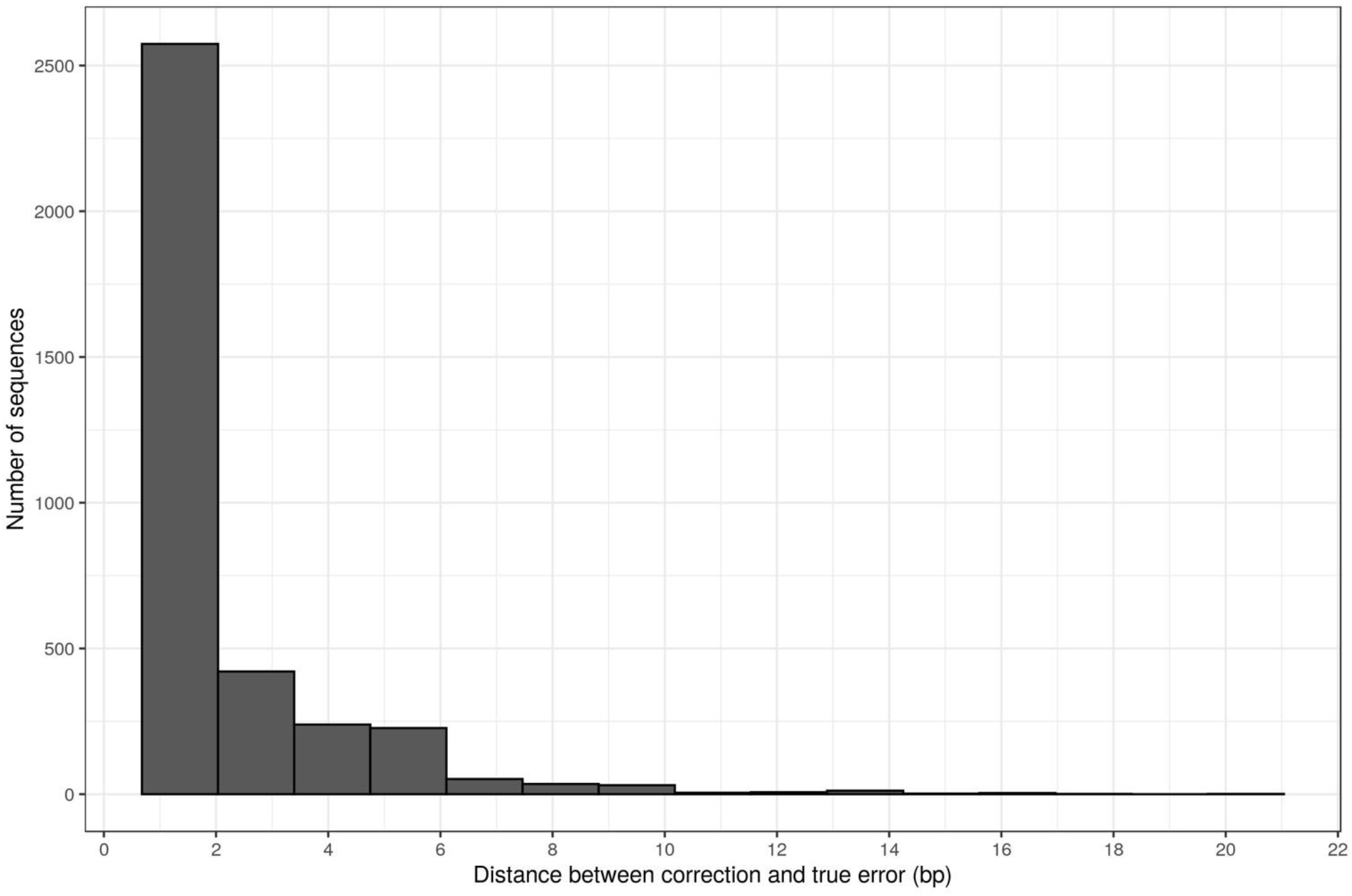
Histogram showing number of base pairs between inexact corrections applied by debar and the ground truth error location for the given sequence. In total 3,612 sequences (36.12%) had errors that were denoised inexactly, and corrections were an average of 2.31 bp (sd = 1.9767) away from the exact ground truth error location.

These results were used to select a default censorship value for debar to ensure that inexactly identified indel errors are masked in most sequences (Figure 1). A default censorship length of 7 (the average miss distance plus two times the standard deviation, rounded up) was selected in order to mask the true error in >95% of instances where inexact corrections were applied, thereby successfully denoising sequences, albeit with some associated loss of information in the sequences, which can be overcome by building a consensus sequence when multiple reads are available for an individual.

Overall, denoising of the 10,000 barcodes with the default censorship parameter (censor_length = 7) resulted in 9,309/10,000 (93.09%) of sequences with errors being successfully denoised. The additional double check parameters (aa_check = True, keep_flanks = False) captured, but did not correct, 269 (2.69%) errors. The debar package thereby corrected or removed 95.74% of sequences with indel errors (Figure 3).

#### False correction rate

A set of 10,000 barcode sequences with no known indel errors was analyzed with debar to determine the incidence of erroneous corrections. Nearly all sequences (99.91%) were not altered nor flagged as erroneous. Nine sequences were erroneously corrected, and none were flagged for rejection. These sequences included a single sequence from each of five orders and four sequences from the order Diptera (flies). Interestingly, the four Diptera sequences that were incorrectly altered all belonged to the same genus: *Culicoides*. They represented 4/58 of all sequences from the family Ceratopogonidae that were in dataset, indicating that the performance issue was isolated to this single genus.

These results indicate that debar deals well with variation in COI sequences across most of the animal kingdom, but that it displays some taxonomic bias in performance. This is a limitation of debar, as any genus with a COI profile that systematically deviates from the COI PHMM used in debar will be erroneously denoised. The benefit of the conservative censorship approach used in the package is that although these reads are erroneously adjusted, the corrections made are masked by Ns, and the entire sequence is not rejected. Rather, only a small section of the sequences is lost, as if it were to contain an indel error. Most of any falsely corrected sequences can thereby be recovered, and in most instances, this would be sufficient to identify associated taxonomy and inform biological conclusions.

#### Denoising PacBio Sequel data

We applied debar in the analysis of real DNA barcode data by developing a processing pipeline (Figure 2 – hereafter ‘the debar pipeline’) and compared the amount of technical noise in the barcodes before and after processing. A set of 29,525 consensus barcode sequences derived from processing data from four Sequel runs were obtained from mBRAVE and were re-processed with the debar pipeline (Table 1).

**Table 1.** Summary of the results for the 29,525 barcode sequences (produced from PacBio Sequel data analyzed using the mBRAVE platform) after processing with the debar pipeline.

| PacBio Sequel run | Run 1 | Run 2 | Run 3 | Run 4 | Total |
| --- | --- | --- | --- | --- | --- |
| Consensus sequences generated | 7,518 | 7,373 | 7,235 | 7,399 | 29,525 |
| Consensus sequences flagged by coil for indel error | 869 | 817 | 900 | 909 | 3,495 (11.8%) |
| Rejected by debar denoising | 8 | 4 | 16 | 9 | 37 (0.1%) |
| Sequences flagged by coil post-denoising | 256 | 285 | 305 | 277 | 1,123 (3.8%) |
| Sequences corrected | 605 | 528 | 579 | 623 | 2,335 (66.8% of flagged sequences) |

Analysis of the consensus barcodes with coil (step ii. of the debar pipeline) flagged 3,495 (11.8% of total) of consensus sequences due to the detection of a stop codon in the translated sequence or due to the presence of an unexpected amino acid (log likelihood score below the default threshold). The large number of flagged sequences is likely reflective of false positives (sequences flagged by coil that lack indel errors due to the incorrect establishment of reading frame). In fact, 2,418 sequences (8.1% of total, 69.2% of flagged sequences) were flagged due to the presence of a stop codon, and 1,282 of them (4.3% of total, 36.7% of flagged sequences) contained a stop codon in all three forward reading frames, providing extremely strong evidence of an indel error (i.e. a low likelihood of being a false positive).

After denoising, the output sequences were again assessed with coil (step viii. of the debar pipeline) and this analysis revealed that debar had corrected many indel errors (Table 1, Table 2). Only 1,123 (3.8%) of the final barcode sequences were flagged by coil’s coi5p_pipe function, suggesting that 66.8% (2,335) of the flagged sequences were successfully denoised. When comparison was restricted to the 2,418 sequences with stop codons, only 176 were still flagged as containing stop codons, indicating that 92.7% (2,242/2,418) of the sequences in this subcategory were effectively denoised. A more conservative estimate of correction success was provided by the subset of flagged sequences with stop codons in all reading frames. Of these sequences, 1106/1282 (86.27%) passed the coil check following denoising, suggesting the successful correction of an indel error and improved representation of the true sequence. External quantification of the debar pipeline’s denoising ability was obtained by the submission of pre- and post- pipeline barcode sequences to BOLD (http://www.boldsystems.org). The sample size for this test was smaller as BOLD requires taxonomic designations and this information was only provided by mBRAVE for 27,041 sequences. The total number of original sequences flagged by BOLD due to its detection of a stop codon was 1,515 (6.3%), a considerably lower frequency than reported by coil on the initial pipeline inputs. Of the 1,515 sequences with initial evidence of stop codons, 14 were rejected outright by the debar pipeline, 223 were flagged but not successfully corrected, 147 were unflagged and not corrected, and 1,131 had no evidence of errors following denoising (Table 3). Based on this assessment with BOLD, the debar pipeline produced a 75% reduction in the number of errors in the dataset from 6.3% (1,515) to 1.6% (384). Of the remaining 384 errors, the majority (223) were detected as problematic and flagged as erroneous by debar. As a consequence, the debar pipeline reduced the number of unidentified errors by >90% (from 1,515 to 147) in the barcode dataset (Table 3).

**Table 2.** Assessment of the correction ability of the debar pipeline for the subset of sequences in the high-confidence error set. This set of sequences was flagged by coil and produced a stop codon when translated within all reading frames. The top half of the table indicates the number of sequences flagged by coil as likely to be erroneous, based on the log likelihood values of the sequences. Results are shown for sequences both before and after the denoising process. The bottom half of the table contains the number of sequences flagged by coil as likely to be erroneous, based on the presence of a stop codon in the amino acid sequence resulting from the censored translation of the framed nucleotide sequence. This high success for the stop-codon metric (86.3% of errors removed) indicates that the pipeline is an effective means of correcting frameshift-causing insertion or deletion errors. The relatively lower success at correcting sequences with low log likelihood values suggests that frameshift-causing errors are not the only set of errors being flagged by coil, and that non-frameshift errors are not effectively corrected by the debar pipeline.

| **PacBio**<br>**Sequel run** | **Run 1** | **Run 2** | **Run 3** | **Run 4** | **Total** |
| --- | --- | --- | --- | --- | --- |
| **Original**<br>**flagged** | 551 | 547 | 609 | 610 | 2,317 |
| **Flagged post-**<br>**denoising** | 254 | 280 | 300 | 271 | 1,105 |
| **Corrected** | 53.9% | 48.8% | 50.7% | 55.6% | 52.3% |
| **PacBio**<br>**Sequel run** | **Run 1** | **Run 2** | **Run 3** | **Run 4** | **Total** |
| **Original stop**<br>**codon** | 319 | 295 | 318 | 350 | 1,212 |
| **Stop codon**<br>**post-**<br>**denoising** | 43 | 42 | 36 | 55 | 176 |
| **Corrected**<br>**stop codons** | 86.5% | 85.7% | 88.7% | 84.2% | 86.3% |

**Table 3.** Result of the BOLD Data System evaluation of debar denoising workflow’s effectiveness. The number of sequences identified by BOLD as containing stop codons, before and after processing with the denoising pipeline (Figure 2). Only the 27,041 specimens with barcodes and taxonomic information produced through the processing of PacBio Sequel data on the MBRAVE platform were considered, as BOLD requires taxonomic information for assessing the presence of stop codons. The rows break the sequences down into categories, which indicate the source of the post-denoising sequence that was submitted to BOLD for assessment.

| Sequence category | Total Sequence count | Stop codon count |  | Percent error reduction |
| --- | --- | --- | --- | --- |
|  |  | Original | Post-denoising |  |
| Unaltered | 23,992 | 88 | 88<sup>†</sup> | - |
| Denoised, altered | 2,265 | 1,190 | 59<sup>†</sup> | 95% |
| Flagged for potential error, unaltered | 701 | 223 | 223* | - |
| Flagged and rejected | 16 | 14 | 14 | - |
| Labelled as *Wolbachia* by MBRAVE | 67 | 0 | 0 | - |
| **Total** | 27,041 | 1,515 (6.3%) | 384 (1.6%) | 74.66% |
| **Total, non-flagged only** | 26,257 | 1,278 (4.8%) | 147 (0.6%) | 88.5% |

<sup>†</sup> The sum of these categories (shown in the final row of the column) represents the false negative rate for the denoising pipeline. These are the 0.6% (147/27,041) of sequences that appear to contain true stop codons that were not flagged for denoising, or that were denoised unsuccessfully and not flagged as potential errors.
\* The false positive rate of the denoising pipeline is the number of sequences in this category that do not in fact contain a stop codon. There is a total of 478 (701-223) false positives and an overall false positive rate of 1.8% (478/27,041). Since this set of sequences are flagged for potential errors, as opposed to being outright rejected, additional inspection of sequences in this category can separate the unsuccessfully denoised sequences with true errors from those that do not contain an error.

The denoising of the barcodes with the debar pipeline did not result in sequences with large amounts of missing information. Of the 29,525 output barcodes, 28,802 were high quality (<1% Ns), 11 were medium quality (<2% Ns), 498 were low quality (<4% Ns), and 214 were unreliable (>4% Ns). There was a strong negative relationship between the number of CCS available for a sample and the amount of missing information in the final barcode sequence (Figure 6).

**Figure 6.**
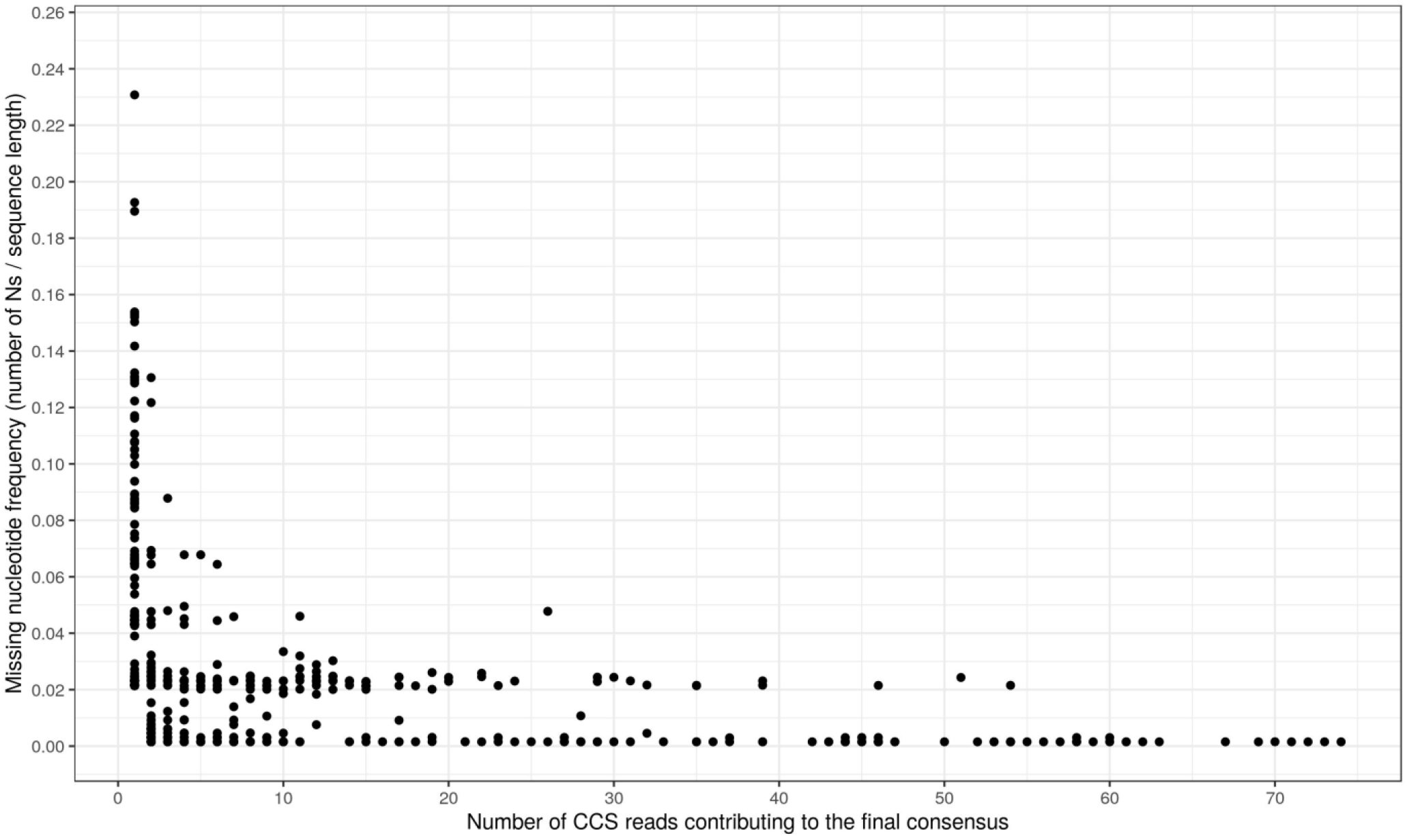
Relationship between the amount of missing data in the final denoised barcode sequences (number of Ns divided by the total length of the sequence) and the number of CCS reads that contributed to the generation of the barcode. The figure displays only the 1,008 denoised barcode sequences submitted to BOLD that contained at least one “N” (the remaining 28,517 barcode sequences in the BOLD submission did not contain an “N”).

### Denoising metabarcode data

#### Consensus sequence quality

Metabarcode data from a mock arthropod community were also denoised followed by comparison of original sequences to the denoised consensus sequences to determine if the debar improved sequence quality (Table 4). Of the original centroid sequences for the 398 BINs, 125/398 (31.4%) contained evidence of indel errors when analyzed with coil. Following denoising and consensus sequence generation via debar, the number of barcode-compliant outputs was considerably higher with only 7/394 (1.8%) displaying evidence of indel errors. Four BINs had all their component sequences rejected by debar so no consensus sequences were generated. The rate of apparent indel errors was higher in the centroids of the 1255 OTUs; 681 (54%) displayed evidence of a stop codon when analyzed with coil, suggesting the presence of indels in more than half of the sequences representing each OTU. The consensus sequences produced through denoising and consensus sequence generation with debar were of apparent higher quality as only 134 (10.6%) displayed evidence of a stop codon when analyzed with coil. An additional 31 OTUs (2.5%) failed to produce a valid consensus sequence after denoising because all their component sequences were rejected by debar.

**Table 4.** Assessment of the sequence quality for data from a mock community of arthropods sequenced in bulk using a Thermo Fisher Ion Torrent and processed on the mBRAVE platform. Sequencing and processing results in two sets of data, groups of sequences assigned to BINs and groups of sequences clustered into OTUs. The representative sequences (centroids before denoising, consensus after denoising) and all individual sequences were checked with the R package coil for evidence of frameshifts (stop codons in amino acid sequence) before and after denoising to see if processing the data with the debar package resulted in higher quality barcode sequences.

| Sequences analyzed | Sequence data source | Original |  | After debar denoising |  |
| --- | --- | --- | --- | --- | --- |
|  |  | Total count | Stop codon count | Total count | Stop codon count |
| Representative sequences | Assigned to BINs | 398 | 125 (31.4%) | 394 | 7 (1.8%) |
|  | OTUs | 1,255 | 681 (54%) | 1,224 | 134 (10.6%) |
| ESVs | Assigned to BINs | 123,926 | 61351 (49.5%) | 122,349 | 2858 (2.3%) |
|  | OTUs | 2,199 | 1310 (59.57%) | 2,145 | 418 (19.49%) |

The corrections did cause some loss of information; 46/394 (11.7%) of the consensus sequences for the BIN groups contained at least one ‘N’ due to ambiguous or censored base pairs in their component reads, and 861/1255 (68.6%) of the OTU consensus sequences contained at least one ‘N’. The number of ‘Ns’ per sequence was generally low for the BINs (median = 0; 12 sequences with 14 or more ‘Ns’) but was higher for the OTUs (median number of ‘Ns’ = 15), indicating there was on average one correction per OTU (correction of an indel, plus the seven bp mask in either direction result in 14 (insertion) or 15 (deletion) consecutive ‘Ns’). There was a positive relationship between the number of sequences within an OTU and the completeness of information in the final consensus sequence.

#### ESV data quality

Data analysis on mBRAVE revealed 398 BINs represented by 123,926 unique dereplicated reads as well as 1255 OTUs lacking taxonomic assignment that were represented by 2199 unique sequence reads. When original sequences were checked with coil, it indicated that 61,351/123,926 (49.5%) of BIN sequences and 1310/2199 (59.97%) of the OTU sequences displayed strong evidence of an indel error as they contained a stop codon when translated. By contrast, following denoising with debar the incidence of stop codons was far lower as just 2858/122,349 (2.3%) of the BIN sequences and 418/2,145 (19.49%) of the OTU sequences had evidence of indels. This result indicated that denoising of individual sequences reduced the incidence of apparent indel errors by over 95% for the BINs (58,593 fewer indel errors) and by 68% for the OTUs (892 fewer indel errors). Most sequences were subjected to at least one indel correction by debar, with 85,298/122,349 (69.7%) of the final BIN sequences and 1387/2145 (64.7%) of final OTU sequences containing at least one ‘N’ character. Low abundance OTUs in the data set represented by biologically valid sequences need not be discarded solely due to their low abundance and could be further inspected for putative evidence of rare community members.

## Discussion

This manuscript introduces debar, a PHMM-based denoiser, and demonstrates how it can improve the quality of sequence data used for both DNA barcode library construction and for metabarcode studies by correcting indels introduced by sequencing error. We first evaluated its effectiveness through an *in silico* study that tested its capacity to recognize and repair reference barcodes with artificially introduced indels. Debar was shown to be effective, as it corrected >95.7% of the errors and applied erroneous adjustments to less than 0.01% of correct sequences. This strong performance extended to real-world data sets. Debar reduced the rate of frameshift indels by 75% in sequence records generated by the long-read Sequel platform, generating more barcode-compliant sequences, most with little or no missing information. Debar also improved the quality of metabarcode data generated by the ION S5 allowing for ESVs to be considered with higher confidence and for the recovery of higher-quality representative sequences for OTUs.

Denoising sequences with artificial errors and known ground truths showed that the corrections performed by debar were imperfect, with the exact indel location being identified only 61.8% of the time. The application of a default 7bp censorship on both sides of putative indel corrections proved to be an effective means of masking most errors, improving the denoiser’s error removal rate to >95.75%. This high error removal rate involves a tradeoff, as sequence adjustments are accompanied with a loss of 14 base pairs of information. This information loss is an acceptable cost, as it ensures that all remaining base pairs can be considered with high confidence. The nature of high-throughput sequence data, namely that there are usually multiple sequencing reads for a given specimen available, can help mitigate the loss of information. Corrected sequences from a specimen or OTU can be used in conjunction with one another, filling in the different censored locations and overcoming the loss of information. The censorship of bases adjacent to indel corrections is an optional parameter that users may alter to suit their needs. Smaller censorship values, or no censorship at all, would result in less loss of information per sequence, but would come at the cost of more errors remaining in the final data.

Denoising of real DNA barcode data obtained from sequencing of specimens on the Pacific Biosciences Sequel platform resulted in higher-quality output sequences. An exact metric quantifying the improvement is, however, difficult to state with certainty, as the ground truth of the sequences is not known. The independent tests of the sequences through submission of consensus sequences to BOLD before and after denoising provided a conservative estimate of the debar package’s effectiveness. Conservatively, this test showed a 75% reduction in the number of barcode sequences with technical indel errors after application of the debar pipeline and a low false negative rate (147 unidentified errors out of 1,515 total putative errors). This is an important improvement because the Pacific Biosciences Sequel platform is used at the Centre for Biodiversity Genomics to produce high-quality reference barcodes for the barcoding research community (Hebert *et al.* 2018). Accuracy of these sequences is therefore important; the debar package is shown to improve sequence quality, yielding more biologically likely and therefore reliable outputs. The generation of barcode sequences is also made more efficient. By increasing the rate of barcode-compliant outputs from 93.7% to 98%, fewer samples require reprocessing or resequencing.

Understanding within-species patterns of genetic diversity is an essential metric for characterizing community health. High intra-species genetic diversity is assumed to indicate healthy ecosystems, comprised of large and stable populations with the standing genetic variation needed to survive environmental stressors (Zizka *et al.* 2020). The characterization of ESVs within OTUs can provide intra-species diversity measures for member species of a community (Frøslev *et al.* 2017). The initial check of the sub-OTU sequence data from the mock community sequenced with IonTorrent revealed a high rate of putative indel errors (54% of sequences), which would lead to a gross over estimation of the number of ESVs within the OTUs. The reduction of the error rate after denoising with debar allows for a more accurate examination of intra-OTU ESVs and therefore allows for more accurate assessments of intra-species diversity and community health, despite the fact that debar is not capable of eliminating non-indel errors from sequences. Even with the improvements to ESV quality by debar, intra-species diversity estimates will likely remain inflated to some extent, as the sequence-by-sequence corrections applied by debar exclusively account for indel errors while substitution errors could persist within the data.

We have demonstrated that debar is an effective means of reducing technical errors in DNA barcode and metabarcode data, but the package is not without limitations. The package is designed to correct insertion and deletion errors, but these are not the only technical issues that can lead to inflated biodiversity estimates. The program is not an effective means of identifying or correcting chimeric sequences or non-animal COI biological contaminants and should these exist within an input data set they are likely to go uncorrected. Additionally, debar does not have the ability to correct substitution errors on a sequence-by-sequence basis. Because of indel correction, denoised sequences are aligned, and nucleotide positions become directly comparable across different sequences from a given specimen or OTU. Random point substitution errors can thereby be corrected in consensus sequence generation, through the ‘majority rule’ approach debar uses in base calling. However, if systematic errors exist (i.e. most sequences possess the same substitution), few sequences are available for consensus sequence generation, or ESVs are being examined, then substitution errors may persist in the data. An additional source of error unaccounted for by debar is contaminant sequences. It has been demonstrated previously that the PHMM utilized in debar is not an effective means of separating animal barcode sequences from off-target barcodes derived from bacteria, plant, fungi, or other origins (Nugent *et al.* 2020). Taken together, these limitations show that debar cannot single handedly address the technical challenges associated with DNA barcoding. The tool is likely most effective when applied in conjunction with existing barcode and metabarcode workflows and improves the quality of final sequences if the inputs have been filtered based on quality, had primers removed, and been cleaned of chimeric and contaminant sequences. The sequence-by-sequence denoising approach of debar means that it is a flexible tool capable of integrating into analysis pipelines for sequencing data from various sources. Application of debar in tandem with conventional, clustering-based denoising tools would likely lead to the highest quality assessment of biodiversity. Following OTU generation with other tools, using debar to denoise all reads within a given OTU prior to consensus sequence generation would maximize accuracy of the consensus sequence while conforming to the conserved structure of the COI barcode region. The removal of intra-OTU noise can also improve the accuracy of alpha-diversity estimates. Additionally, application of debar in the denoising of rare, low-abundance sequences not present in the OTUs would allow these data to be further examined with higher confidence, revealing biological insights that would be overlooked in conventional workflows.

The PHMM denoising technique used by debar is an effective barcode-focused framework that can be extended to fit a variety of needs. Data from only two sequencing platforms were tested in this study: the Pacific Biosciences Sequel and Thermo IonTorrent S5. Since the PHMM used in debar is barcode specific and not sequencer specific, debar can be effectively applied in denoising of barcode data obtained from any sequencing platform. However, the effectiveness of the denoiser will depend on the types and rates of technical errors associated with a given platform. When applied to data from sequencers such as the Illumina MiSeq, the rate of technical errors corrected by debar will be lower, as this platform is more prone to introduction of substitution, as opposed to indel, errors (Schirmer *et al.* 2015). Although the debar package contains a PHMM for only the common animal barcode COI, the denoising algorithm can in the future be extended and applied in the correction of data for other DNA barcodes with conserved structures.

## Conclusion

This study has described debar, an R package for denoising DNA barcode data, and demonstrated its ability to correct indels in both barcode and metabarcode sequences due to instrument error. In each dataset, debar improved sequence quality. It reduced the apparent number of indels by 75% in data generated by Sequel, increasing the proportion of sequences that met the quality standards required to qualify as a reference barcode. The merits of debar for metabarcode analysis were twofold, allowing more likely consensus sequences to be obtained for OTUs, and for intra-OTU variation to be quantified with higher confidence. Overall, debar is a robust utility for identifying deviations from the highly conserved protein-coding sequence of the COI barcode region. Corrections informed by its use improve the separation of true biological variation from technical noise, with low frequencies of false corrections. Integration of debar into the workflows for processing barcode and metabarcode data will allow biological variation to be characterized with higher accuracy.

## Supporting information

debar_supplementary

## Acknowledgements

This research was supported by grants from Genome Canada through Ontario Genomics and from the Ontario Ministry of Economic Development, Job Creation and Trade. The funders played no role in study design or decision to publish. This research was enabled in part by resources provided by Compute Canada (www.computecanada.ca). We thank Tony Kuo and Thomas Braukmann for aid with data acquisition and interpretation and Tony for helpful comments on the manuscript.

## Data Accessibility Statement

DNA barcode sequences used in training of the Profile Hidden Markov Models are available in the Supplementary data of the following paper: https://doi.org/10.1139/gen-2019-0206. DNA barcode sequences used in model testing are available in this manuscript’s Supplementary files. The R source code for the debar package is available on GitHub: https://github.com/CNuge/debar. Additional data and code available on request from the authors.

## Author Contributions

The study was conceived and designed by SJA, PDNH, SR, and CMN. The programming of the debar package was performed by CMN. Analyses of package performance were performed by CMN with resources, design, and other assistance provided by TAE, SR, and SJA. The initial draft of the manuscript was written by CMN and SJA. All authors contributed to the editing of the manuscript.

## Supplementary Information

**Supplementary File 1** (‘S1-single_errors_in_10k_sequences.csv’) The 10,000 COI barcode sequences with single introduced indel errors that were used to test debar and calibrate the default parameters.

**Supplementary File 2** (‘S2-control_denoising_no_errors.csv’) The 10,000 COI barcode sequences with no known indel errors used to assess the false correction rate of debar

**Supplementary File 3** (‘S3-single_file_pipeline’) Scripts and example data for the denoising pipeline developed to process COI DNA barcode sequence data produced using the Pacific BioSciences Sequel sequencer and mBRAVE platform

**Supplementary File 4** Scripts and example data for the denoising pipeline developed to process COI DNA metabarcode sequence data produced using the IonTorrent S5 sequencer and the mBRAVE platform

**Supplementary File 5** Vignette demonstrating the functionality of the debar package. The vignette is also available as part of the R package (https://github.com/CNuge/debar/tree/master/vignettes)

