## Supplementary material for "debar, a sequence-by-sequence denoiser for COI-5P DNA barcode data": debar_supplementary: consensus_selection.r

```
################
# take the denoising outputs, select and label the optimal set of consensus sequences
################

# once the denoise_pipeline.r runs successfully and generates all outputs, then take the code 
# from generate_output_seqs.r and use it to generate the single file optimal outputs file
# will need to read in the new equivalents of the old files - so compare denoise_pipeline.r
# to the write locations for its predecessors from:
# assess_improvement, eval_denoised_components, and denoise_components

library(tidyverse)
library(parallel)

source('R/sequel_processing_functions.r')


main = function(original_coil_outputs, denosed_consensus, denoised_coil_check,
		drop_control = TRUE){

	print("reading in the data components")
	###
	#1 original consensus sequences evaluated by coil
	print("1-original consensus sequences evaluated by coil")
	original_consensus_coil_dat = read_tsv(original_coil_outputs)
	original_consensus_coil_dat$mid_id = unlist(lapply(original_consensus_coil_dat$name, 
	function(x){
		strsplit(x, "\\|")[[1]][[1]]												
	}))

	#drop the control sequences - not needed
	if (drop_control == TRUE){
		original_consensus_coil_dat = original_consensus_coil_dat[!grepl("CONTROL",original_consensus_coil_dat$mid_id),]
	}

	###
	#2 denoiser sequences
	print("2-denoised sequences")
	denoised_out_dat = read_tsv(denosed_consensus)

	#add a column with rank_mid so that the df can be merged easily after coil analysis.
	denoised_out_dat$mid_id_rank = unlist(lapply(1:nrow(denoised_out_dat), function(i){
		paste0(denoised_out_dat$rank[[i]], "_", denoised_out_dat$mid_id[[i]])
		}))

	###
	#3 denoiser output coil checks
	print("3-denoised outputs checked with coil")
	denoised_coil_check = read_tsv(denoised_coil_check)
	#split the name components
	denoised_coil_check$rank = unlist(lapply(denoised_coil_check$name, function(x){
		as.integer(strsplit(x, '\\_')[[1]][[1]])
		}))

	denoised_coil_check$mid_id = unlist(lapply(denoised_coil_check$name, function(x){
		strsplit(x, '\\_')[[1]][[2]]
		}))

	denoised_coil_check = denoised_coil_check[!grepl("CONTROL",denoised_coil_check$mid_id),]

	write_optimal_out(original_consensus_coil_dat, denoised_out_dat, denoised_coil_check)
}


#the original consensus sequences
original_coil_outputs = 'pipeline_outputs/coil_evaluated.tsv'

#the denoised consensus sequences, and their coil check values
denosed_consensus = 'pipeline_outputs/denoised_reads.tsv'
denoised_coil_check = 'pipeline_outputs/denoised_consensus_coil_outputs.tsv'


main(original_coil_outputs, denosed_consensus, denoised_coil_check)
```
