## Supplementary material for "debar, a sequence-by-sequence denoiser for COI-5P DNA barcode data": debar_supplementary: denoise_pipeline.r

```
################
# take in the mBRAVE consensus sequence list, and the cleaned component data.
# evaluate consensus sequences, denoise and regerate consensus where needed.
################

library(coil)
library(debar)
library(tidyverse)
library(parallel)
library(kmer)

source('R/debar_pipeline_functions.r')


#coil 
main  = function(consensus_filename, cleaned_reads_filename, 
					ncores=1, censor_length=7, outdir = 'pipeline_outputs/',
					coil_check == TRUE){

	dir.create(outdir)
	
	#output filenames  
	#coil_1_outfile = paste0(outdir, 'coil_evaluated_', gsub(".fas",".csv",consensus_filename))
	coil_1_outfile = paste0(outdir, 'coil_evaluated.tsv')
	#denoised_metadata_file = 
	denoised_metadata_file = paste0(outdir, 'denoising_metadata.tsv')
	denoised_consensus_outfile = paste0(outdir, 'denoised_reads.tsv')

	print("reading data")
	data = read_fasta(consensus_filename)

	print("parsing header")
	data2 = do.call(rbind, lapply(data$header, function(x){
		parse_consensus_header(x)
		}))

	data = cbind(data, data2)

	for(i in names(data)){
		data[[i]] = as.character(data[[i]])
	}

	data = as_tibble(data, stringsAsFactors=FALSE)

	print("running coil analysis")
	coil_data = mclapply(1:nrow(data), function(i){
		coi5p_pipe(data$sequence[[i]], 
				name = data$header_data[[i]],
				indel_threshold = -358.88)
		}, mc.cores = ncores)


	print("flattening coil output")
	out_data = flatten_coi5p(coil_data)
	out_data = as_tibble(out_data)
	out_data$MID = unlist(lapply(out_data$name, function(x){
		mid_extract(x)
		}))

	print("writing coil data to file")
	write_tsv(out_data, coil_1_outfile)

	print("subsetting error sequences")
	flagged = out_data[out_data$indel_likely == TRUE | out_data$stop_codons == TRUE ,]


	# TODO - decide if you want to output this, in addition to, or instead of 
	# not_flagged = data[data$indel_likely != TRUE & data$stop_codons != TRUE ,]

	print("loading cleaned ccs data file")
	ccs_data = read_tsv(cleaned_reads_filename)

	print("subsetting reads for problem consensus")
	ccs_data = ccs_data[ccs_data$MID %in% flagged$MID,]

	print("denoising required reads")
	#to get full multicore benefit, probably best to denoise here
	ccs_data$denoised_seq = denoise_list(ccs_data$trimmed_seq, dir_check = FALSE,
												keep_flanks = 'right',
												censor_length = censor_length,
												cores = ncores)


	print("identifying rejections")
	#turn nulls to NAs, then unlist to make df column
	ccs_data$denoised_seq = unlist(lapply(ccs_data$denoised_seq, function(x){
			if(is.null(x)){
				return(NA)
			}else{
				return(x)
			}}))

	print("splitting on mid tags")
	#split on the sample id
	data_mids = split(ccs_data, ccs_data$MID)

	print("number of mid tags:")
	print(length(data_mids))

	print("generating metadata")
	met_data = data.frame(mids = names(data_mids),
							org_seq_count = unlist(lapply(data_mids, nrow)),
							rejected_reads = unlist(lapply(1:length(data_mids), function(x){
								sum(is.na(data_mids[[x]]['denoised_seq']))
								})),
							stringsAsFactors=FALSE)	

	print("generating otus and consensus sequences")
	den_outputs = mclapply(1:length(data_mids), function(i){
			mid_id = names(data_mids[i])
			
			#denoise all of the sequences in the list
			#seqs = data_mids[[i]][['trimmed_seq']]
			#denoised_list = denoise_list(seqs, dir_check = FALSE,
			#										keep_flanks = 'right',
			#										cores = parallel::detectCores())
			denoised_list = data_mids[[i]][['denoised_seq']]

			#switch this comment to following line if you only want the top sequence
			#top_c_out = top_otu_consensus(denoised_list)
			all_c_out = all_otu_consensus_seqs(denoised_list)

			all_c_out$mid_id = mid_id

			return(all_c_out)
		}, mc.cores = parallel::detectCores())	

	print("building full OTU dataframe")
	den_out_df = do.call("rbind", den_outputs)

	print("writing denoised data to file")
	write_tsv(den_out_df, denoised_consensus_outfile)
	write_tsv(met_data, denoised_metadata_file)

	if(coil_check == TRUE){
		denoise_coil_check(denoised_consensus_outfile)
	}

}


args = commandArgs(trailingOnly = TRUE)

for(i in 1:length(args)){
	a = args[[i]]
	if((a == '--consensus') | (a == '-c')){
		consensus_filename = args[[i+1]]
	}else if((a == '--read_data') | (a == '-r')){
		cleaned_reads_filename = args[[i+1]]
	}
}

print("arguments passed:")
print(paste0("consensus: ", consensus_filename))
print(paste0("read_data: ", cleaned_reads_filename))

#consensus_filename = '../consensus_seqs/342_SMRT-00289.08-11.41385.consensusseq.fas' #this needs to be .fasta
#cleaned_reads_filename = 'component_data_cleaned_41385.tsv'

ncores = detectCores()
main(consensus_filename, cleaned_reads_filename, ncores)

#Full pipeline tested - its working!
```
