## Supplementary material for "debar, a sequence-by-sequence denoiser for COI-5P DNA barcode data": debar_supplementary: debar_pipeline_functions.r

```
parse_consensus_header = function(header){

	header_names = c('id', 'run', 'contig_id', 'rep_count', 'id_similarity', 
						'c_count', 'cmxd', 'cmnd', 'cnnd', 'p', 'c', 'o', 
						'f', 'g', 's', 'otu', 'date')

	header = gsub("otu:BOLD:", "otu:", header)

	#split the header on the two delimiters, take the non-names
	split_header = unlist(strsplit(header, '[|:]'))[c(TRUE, FALSE)]
	
	#label the list
	names(split_header) = header_names

	return(split_header)

}

mid_extract = function(x){
	y = strsplit(x, "\\|")[[1]][[1]]
	return(y)
}


#' Determining the most common character in each column.
#'
#' @keywords internal
col_mode = function(col_data) {
  if (!is.factor(col_data)){
    col_data <- factor(col_data)
  } 
  
  tab_col = tabulate(col_data)
  
  if(max(tab_col) == 0){
    return("n")
  }
  
  consensus = levels(col_data)[tab_col == max(tab_col)]
  
  if (length(consensus)>1) {
    return("n")
  } 
  
  return(consensus)
}

#' build an DNAbin with ape.
#'
#' Switches the dashes in the seq - to n
#'
#' @keywords internal
list_to_DNAbin = function(dna_strings){
	return(ape::as.DNAbin(strsplit(gsub('-', 'n', as.character(tolower(dna_strings))),"")))
}

#rejects the long sequence and then produces new outputs,
#these can be combined with consensus_sequence to make
#a sequence with not stop codons in the AA, where all are in frame.

#check - if any are out of phase, or if there is weird behaviour with the
#consensus, there appears to be a problem on the combination step, i.e.
# things that are too dissimilar from one another are getting combined,
# missing values aren't being ignored properly or something else to this effect.

#either - add a cluster step post denoising prior to consenus, generating multiple
#barcodes for a sample (need to record the #) or pump the reads
#to an output and plug into the current mBRAVE consensus algorithm.


top_otu_consensus = function(denoised_list, threshold = 0.975){
	
	denoised_list = unlist(denoised_list)
	denoised_list=denoised_list[!is.na(denoised_list)]

	x = otu(list_to_DNAbin(denoised_list) , threshold = threshold)

	top_otu = denoised_list[x == col_mode(x)]

	con = consensus_sequence(top_otu)
	
	return(con)

}


#' Take the denoised list of sequences from Debar, use the kmer package to 
#' cluster sequences into OTUs and return the consensus sequence for each OTU
all_otu_consensus_seqs = function(denoised_list, threshold = 0.975){

	denoised_list = unlist(denoised_list)
	denoised_list = denoised_list[!is.na(denoised_list)]

	if(length(denoised_list)==0){
		return(data.frame(consensus = '', 
					rank = 1,
					count = 0, 
					stringsAsFactors=FALSE))
	}

	otus = otu(list_to_DNAbin(denoised_list), threshold = threshold)
  
	tab_col = table(otus)
	
	tab_keep = sort(tab_col, decreasing = TRUE)

	consensus = c()
	rank = c()
	count = c()


	#If there are reads that met the frequency levels
	for(i in 1:length(tab_keep)){
		g = names(tab_keep)[[i]]
		members = denoised_list[otus == g]
		con = consensus_sequence(members)
		
		consensus = c(consensus, con)
		rank =c(rank, i) 
		count = c(count, tab_keep[[i]])
	}

	out_df = data.frame(consensus = consensus, 
					rank = rank,
					count = count, 
					stringsAsFactors=FALSE)

	return(out_df)
}


denoise_coil_check = function(denosed_filename){

	#read in the denoised reads
	denoised_out_dat = read_tsv(denosed_filename)

	print("building mid_rank column")
	#add a column with rank_mid so that the df can be merged easily after coil analysis.
	denoised_out_dat$mid_id_rank = unlist(lapply(1:nrow(denoised_out_dat), function(i){
		paste0(denoised_out_dat$rank[[i]], "_", denoised_out_dat$mid_id[[i]])
		}))

	print("running coil on denoised outputs")
	denoised_consensus_coil_output = list()
	print("dropping NA sequences (failed debar) prior to coil check")
	#drop the na sequences that were rejected by debar
	denoised_out_dat = denoised_out_dat[!is.na(denoised_out_dat$consensus),]
	print("running coil check")
	#build the denoiser outputs
	denoised_consensus_coil_output = mclapply(1:nrow(denoised_out_dat), function(i){
			coi5p_pipe(denoised_out_dat$consensus[[i]], 
					name = denoised_out_dat$mid_id_rank[[i]],
					trans_table = 0,
					indel_threshold = -358.88)
		}, mc.cores = detectCores())

	print("flattening coil output")
	#flatten this, merge back with the denoised_out_dat
	denoised_coil_df = flatten_coi5p(denoised_consensus_coil_output)
	denoised_coil_df = as_tibble(denoised_coil_df)


	print("writing denoised coil check to file")
	outname = paste0("pipeline_outputs/denoised_consensus_coil_outputs.tsv")
	write_tsv(denoised_coil_df,outname)

}

s2v = function(x, lowercase = TRUE){
  strsplit(x, "")[[1]]
}
v2s = function(x){
  paste(x, collapse = "")
}

raw_str = "nnnntatattttatctttggcgcatgagcagggatactaggtacatccttaagaatcctcatccgaacagaattaggtcaaccaggttctctaattggtgacgatcaaatttataatgtaatcgttacagcacgtgccttcatcataattttcttcatagttatgcctattataatcggtggttttggaaattgactagtacctttaatacttggagcacctgatatagcattcccacgaataaataacataagattttgattattaccaccatcaatccttctccttctaattagaagaactgttgaaagtggggcgggtacaggatgaactgtatatccccctctttcagctagaattgcccatgctgggcctgcagtagatttaactattttttcacttcatcttgctgggatatcaagaatcataggtgcagtaaattttattacaacaataattaacataaaaccaatttatataaaccaaacacaaataccactatttgtttgatcagttggtactactgctcttttacttttattatctcttcctgtattagcgggggctattactatactattaacagatcgtaatctaaatacatcttttttcgaccctgcaggtgtgggggatccaattctataccaacactt"

# a function to take the leading ns of of an DNA sequence
leadn_strip = function(raw_str){
	raw_vec = s2v(toupper(raw_str))
	for(i in 1:length(raw_vec)){
		if(raw_vec[[i]] != "N"){
			return(v2s(raw_vec[i:length(raw_vec)]))
		}
	}
	warning("passed sequence is all N characters")
	return(raw_str)
}

# take the different data sources (original coil check, denoised sequences and denoised coil data)
# and generate the optimal set of output sequences
write_optimal_out = function(original_consensus_coil_dat, denoised_out_dat, denoised_coil_check){
	print("building output")
	output_sequences = tibble(mid_id = character(),					#the mid_id
								original_sequence = character(),	#the original output sequence
								passed_initial_coil = logical(),	#inital coil pass val
								initial_aaScore = numeric(),		#inital aa logl score
								initial_stop_codons = logical(),	#were there stop codons
								selected_sequence = character(),	#the final output sequence
								selected_source=character(),		#either original,denoised_1 or denoised_2
								selected_aaScore = numeric(),
								selected_stop_codons = logical(),		#are the NNNNNNN sections in the output
								n_count = numeric(),
								original_header = character(),
								stringsAsFactors=FALSE)				#how many Ns are in the sequence

	print("gathering sequences")
	#iterate through all the sequences in the initial coil output dataframe
	for(i in 1:length(original_consensus_coil_dat$mid_id)){
		seq = original_consensus_coil_dat$mid_id[[i]]
		initial_out = original_consensus_coil_dat[original_consensus_coil_dat$mid_id == seq,]

		if(initial_out$indel_likely == FALSE & initial_out$stop_codons == FALSE){

			outseq = toupper(initial_out$raw)
			new_seq = list(mid_id = seq,					
				original_sequence = outseq,	
				passed_initial_coil = TRUE,	
				initial_aaScore = initial_out$aaScore,
				initial_stop_codons = initial_out$stop_codons,
				selected_sequence = outseq,
				selected_source = "initial",
				selected_aaScore = initial_out$aaScore,
				selected_stop_codons = initial_out$stop_codons,
				n_count = lengths(regmatches(outseq, gregexpr("N", outseq))),
				original_header = initial_out$name)

		}else{
			denoised_seqs = denoised_coil_check[denoised_coil_check$mid_id == seq,]
			top = denoised_seqs[denoised_seqs$rank == 1,]

			if(nrow(top) == 0){
				outseq = toupper(initial_out$raw)
				new_seq = list(mid_id = seq,					
					original_sequence = outseq,	
					passed_initial_coil = FALSE,	
					initial_aaScore = initial_out$aaScore,
					initial_stop_codons = initial_out$stop_codons,
					selected_sequence = outseq,
					selected_source = "all_rejected",
					selected_aaScore = initial_out$aaScore,
					selected_stop_codons = initial_out$stop_codons,
					n_count = lengths(regmatches(outseq, gregexpr("N", outseq))),
					original_header = initial_out$name)
				
			}else if(top$indel_likely == FALSE & top$stop_codons == FALSE){
				outseq = toupper(top$raw)
				outseq = leadn_strip(outseq)
				
				new_seq = list(mid_id = seq,					
					original_sequence = toupper(initial_out$raw),	
					passed_initial_coil = FALSE,	
					initial_aaScore = initial_out$aaScore,
					initial_stop_codons = initial_out$stop_codons,
					selected_sequence = outseq,
					selected_source = "denoised_1",
					selected_aaScore = top$aaScore,
					selected_stop_codons = top$stop_codons,
					n_count = lengths(regmatches(outseq, gregexpr("N", outseq))),
					original_header = initial_out$name)

			}else if(nrow(denoised_seqs) > 1){
				if(denoised_seqs$stop_codons[[2]] == FALSE & denoised_seqs$indel_likely[[2]] == FALSE){
					second = denoised_seqs[denoised_seqs$rank == 2,]
					outseq = toupper(second$raw)

					new_seq = list(mid_id = seq,					
						original_sequence = toupper(initial_out$raw),	
						passed_initial_coil = FALSE,	
						initial_aaScore = initial_out$aaScore,
						initial_stop_codons = initial_out$stop_codons,
						selected_sequence = outseq,
						selected_source = "denoised_2",
						selected_aaScore = second$aaScore,
						selected_stop_codons = second$stop_codons,
						n_count = lengths(regmatches(outseq, gregexpr("N", outseq))),
						original_header = initial_out$name)
				
				}else{
					outseq = toupper(initial_out$raw)
					new_seq = list(mid_id = seq,					
						original_sequence = outseq,	
						passed_initial_coil = FALSE,	
						initial_aaScore = initial_out$aaScore,
						initial_stop_codons = initial_out$stop_codons,
						selected_sequence = outseq,
						selected_source = "initial_error",
						selected_aaScore = initial_out$aaScore,
						selected_stop_codons = initial_out$stop_codons,
						n_count = lengths(regmatches(outseq, gregexpr("N", outseq))),
						original_header = initial_out$name)	
				
				}
			}else{
				outseq = toupper(initial_out$raw)
				new_seq = list(mid_id = seq,					
					original_sequence = outseq,	
					passed_initial_coil = FALSE,	
					initial_aaScore = initial_out$aaScore,
					initial_stop_codons = initial_out$stop_codons,
					selected_sequence = outseq,
					selected_source = "initial_error",
					selected_aaScore = initial_out$aaScore,
					selected_stop_codons = initial_out$stop_codons,
					n_count = lengths(regmatches(outseq, gregexpr("N", outseq))),
					original_header = initial_out$name)	
			}
		}
		#append the new output row to the dataframe
		output_sequences = rbind(output_sequences, new_seq, stringsAsFactors=FALSE)
	}

	#add column indicating if there are Ns in the entry
	output_sequences$has_n = unlist(lapply(output_sequences$n_count, function(x){
		if(x == 0){
			return(FALSE)
		}
		return(TRUE)
		}))

	print("writing to file")
	outfile = paste0("pipeline_outputs/final_output_sequences.tsv")
	write_tsv(output_sequences, outfile)
}
```
