## Supplementary material for "debar, a sequence-by-sequence denoiser for COI-5P DNA barcode data": debar_supplementary: sequel_processing_functions.r

```
library(seqinr)
library(tidyverse)
library(janitor)
library(parallel)

######################
#
# functions 
#
######################

#' get the reverse compliment of a DNA sequence read
rev_comp = function(x){
  return(seqinr::c2s(rev(seqinr::comp(seqinr::s2c(x)))))
}

#' Trim the raw ccs read based on the information provided
trim_sequence = function(sequence, orientation, 
						f_pad_len = 5,  
						f_mid_len = 16,
						f_adapter_len = 30, 
						f_primer_len = 30,
						r_pad_len = 5,  
						r_mid_len = 16,
						r_adapter_len = 30, 
						r_primer_len = 30,
						f_start = 1,
						f_mid_ins = 0,
						f_mid_dels = 0,
						r_start = 1,
						r_mid_ins = 0,
						r_mid_dels = 0
						){

	if(orientation == "RC"){
		sequence = rev_comp(sequence)
	}

	#account for length of the mids
	
	f_mid_len = f_mid_len - f_mid_ins + f_mid_dels
	r_mid_len = r_mid_len - r_mid_ins + r_mid_dels
	
	#sum the lengths of the components
	
	f_trim = f_pad_len + f_mid_len + f_adapter_len + f_primer_len + 1 - f_start
	r_trim = r_pad_len + r_mid_len + r_adapter_len + r_primer_len + 1 - r_start
		

	seq_vec = strsplit(sequence, '')[[1]]	
	trimmed_seq	= seq_vec[(f_trim+1) : (length(seq_vec)-r_trim)]

	return( paste(trimmed_seq, collapse = "") )
}


mid_tag_pipe = function(seq_dat, met_dat){

	#subset the met data for only the mid tags in the consensus seq list
	met_dat = met_dat[!is.na(met_dat$MID),]

	#subset the reads in the remaining meta data
	#
	seq_dat$READ = unlist(lapply(seq_dat$header_data, function(x){
		strsplit(x, '\\|')[[1]][[1]]
	}))

	seq_dat = seq_dat[seq_dat$READ %in% met_dat$READ, ]

	seq_dat$MID = unlist(lapply(seq_dat$header_data, function(x){
		strsplit(x, 'MID=')[[1]][[2]]
	}))

	#each read has a meta data component?
	#nrow(seq_dat) == nrow(met_dat)
	
	#make them tibbles
	seq_dat = as_tibble(seq_dat)
	met_dat = as_tibble(met_dat)

	#join the two dataframes.
	read_data = full_join(met_dat, seq_dat, by = c('READ', "MID"))
	#set the name style so its all the same, and matches the mBRAVE style
	read_data = clean_names(read_data, "screaming_snake")

	read_data$raw_lengths = unlist(lapply(read_data$SEQUENCE, nchar)) 
	#apply the trimming to each of the sequences
	#this function accounts for the amount of missing info and any indels in the mid tags
	#takes the reverse compliment of the read when necessary
	read_data$trimmed_seq = unlist(mclapply(1:nrow(read_data), function(i){
		trim_sequence(sequence = read_data[["SEQUENCE"]][[i]],
						orientation = read_data[["ORIENTATION"]][[i]],
						f_start = read_data[["X5P_START"]][[i]],
						f_mid_ins = read_data[["X5P_INS"]][[i]],
						f_mid_dels = read_data[["X5P_DEL"]][[i]],
						r_start = read_data[["X3P_START"]][[i]],
						r_mid_ins = read_data[["X3P_INS"]][[i]],
						r_mid_dels = read_data[["X3P_DEL"]][[i]])
	}, mc.cores = detectCores()))

	read_data$trimmed_seq = tolower(read_data$trimmed_seq)

	read_data$trimmed_lengths = unlist(lapply(read_data$trimmed_seq, nchar)) 

	return(read_data)	
}
```
