## Supplementary material for "debar, a sequence-by-sequence denoiser for COI-5P DNA barcode data": debar_supplementary: README.md

# debar pipeline

A processing workflow for a single PacBio sequel run from mBRAVE.

1. `sequel_json_processing.r`
	- take in the raw fq and json data from mbrave
	- clean and combine the two data sources
	- write to specified output tsv

*example command:*
```
Rscript sequel_json_processing.r --fq ./ex_data/08-11.41385-demux.0000.fastq.gz --json ./ex_data/08-11.41385-demux_mapping.0000.json.gz --outfile component_data_cleaned_41385.tsv

```

2. `denoise_pipeline.r`
	- take in the consensus sequences (fasta format) from mBRAVE and the output from step 1
	- check sequences with coil, grab component data for failures and denoise, regerate consensus sequences, optionally check new consensus sequences with coil
	- all outputs written to the pipeline_outputs folder
*example command:*
```
Rscript denoise_pipeline.r --consensus ./ex_data/342_SMRT-00289.08-11.41385.consensusseq.fas --read_data component_data_cleaned_41385.tsv

```

3. `consensus_selection.r`
	- take in the set of 3 outputs generated by step 2, select the optimal set of sequences for final denoised output. 
	- generates a tsv with the optimal outputs and records the data sources
	- This one relies on following the denoising pipeline, so assumes the following files:
		- pipeline_outputs/coil_evaluated.tsv
		- pipeline_outputs/denoised_reads.tsv
		- pipeline_outputs/denoised_consensus_coil_outputs.tsv


*example command:*
```
Rscript consensus_selection.r
```


- can wrap all three of the above in a shell script that calls them sequentially, just need the 4 unique filenames provided.
