## Supplementary material for "debar, a sequence-by-sequence denoiser for COI-5P DNA barcode data": debar_supplementary: sequel_json_processing.r

```
#############
# take in a sequel fasta file, and the 
#############

library(debar)
library(janitor)
library(jsonlite)
library(parallel)
library(seqinr)
library(tidyverse)

source('R/sequel_processing_functions.r')


main  = function(fq_inpt, metadata_input, outfile){

	# read in the sequel fastq data
	seq_dat = read_fastq(fq_input)
	# read in the mBRAVE meta data for the sequences
	met_dat_json = fromJSON(metadata_input)
	#build df from relevant component
	met_dat = as_tibble(met_dat_json[['metadata']])

	processed_data = mid_tag_pipe(seq_dat, met_dat)

	#write_the data to file
	write_tsv(processed_data, outfile)

}


# command line argument parsing to get the relevant filenames
args = commandArgs(trailingOnly = TRUE)

for(i in 1:length(args)){
	a = args[[i]]
	if((a == '--fq') | (a == '-f')){
		fq_input = args[[i+1]]
	}else if((a == '--json') | (a == '-j')){
		metadata_input = args[[i+1]]
	}else if((a == '--outfile') | (a == '-o')){
		outfile = args[[i+1]]
	}
}

print("arguments passed:")
print(paste0("fastq: ", fq_input))
print(paste0("json: ", metadata_input))
print(paste0("outfile: ", outfile))

#uncomment these and change string to control directly from within R session
#fq_input = '../sequel_data/08-11.41385-demux.0000.fastq.gz'
#metadata_input = '../sequel_data/08-11.41385-demux_mapping.0000.json.gz'
#outfile = "component_data_cleaned_41385.tsv" 

print("processing data from files")
main(fq_input, metadata_input, outfile)

print("done")
```
