## Supplementary material for "debar, a sequence-by-sequence denoiser for COI-5P DNA barcode data": debar_supplementary: cluster_analysis.r

```
library(tidyverse)
library(kmer)
library(coil)
library(debar)
library(parallel)

list_to_DNAbin = function(dna_strings, ids = NULL){

	x = strsplit(gsub('-', 'n', as.character(tolower(dna_strings))),"")
	
	if(!is.null(ids)){
		names(x) = ids
	}
	
	bin = ape::as.DNAbin(x)

	return(bin)
}

is_rep = function(x){
	return(grepl("\\*$", x))
}

get_otu_centroid = function(otu_df){
	clust_dat = otu(list_to_DNAbin(otu_df$seq), k = 4, threshold = 0.95, method = 'centroid')

	is_cent = unlist(lapply(names(clust_dat), is_rep))

	otu_df$centroid = unlist(lapply(1:length(clust_dat), function(i){
		clust_dat[[i]] == 1 & is_cent[[i]]
		}))

	return(otu_df)
}

#############
# read in the data

original_data = read_tsv('data/denoised_run_18333_derep_data_keepright.tsv')
original_data #129,564 sequences
#128,767 sequences not NA in the denoise col

#####
# subset the data for the OTU data
den_dat = original_data[c('label', 'denoised_seqs')]
den_dat

####
# MBRAVE OTU data
mbrave_otu = read_tsv('data/mBRAVE_OTUs.tsv')
mbrave_otu

#############
# number of OTUs

length(unique(mbrave_otu$otu)) # 5102 
otu_components = table(mbrave_otu$otu)

##############
# have lavels BOLD: chimera or OTU
# pull the BOLD - see if number is 1255
# pull the OTU - see what these are, low coverage numbers?

good_clus_seqs = mbrave_otu[grepl('BOLD', mbrave_otu$otu),]
mis_clus_seqs = mbrave_otu[grepl('TAX', mbrave_otu$otu),]
chimera_clus_seqs = mbrave_otu[grepl('chimera', mbrave_otu$otu),]
OTU_clus_seqs = mbrave_otu[grepl('OTU', mbrave_otu$otu),]

nrow(good_clus_seqs)
nrow(mis_clus_seqs)
nrow(chimera_clus_seqs)
nrow(OTU_clus_seqs)


##############
# join the good OTUs with the denoised sequenes
named_otu_data = rbind(good_clus_seqs, mis_clus_seqs)
named_otu_data = left_join(named_otu_data, den_dat)
named_otu_data


table(named_otu_data$otu)
max(table(named_otu_data$otu)) 
min(table(named_otu_data$otu)) 
mean(table(named_otu_data$otu)) #311.37
sd(table(named_otu_data$otu)) #311.37
median(table(named_otu_data$otu)) #110.5
hist(table(named_otu_data$otu))

# join the unnamed OTUs with the denoised sequenes
unnamed_otu_data = left_join(OTU_clus_seqs, den_dat)

#get the OTU labels for the unnamed OTUs
unnamed_otu_data$original_centroid = unnamed_otu_data$otu_centroid == unnamed_otu_data$label
#ensure one rep per centroid
length(unique(unnamed_otu_data$otu)) == sum(unnamed_otu_data$original_centroid)

write_tsv(named_otu_data, 'data/named_bins_data.tsv')
write_tsv(unnamed_otu_data, 'data/all_otu_data.tsv')


split_named_otus = split(named_otu_data, named_otu_data$otu)
split_unnamed_otus = split(unnamed_otu_data, unnamed_otu_data$otu)


#get the centroid labels for the named OTUs
split_named_otus = lapply(split_named_otus, get_otu_centroid)
split_unnamed_otus = lapply(split_unnamed_otus, get_otu_centroid)


##################
# coil check wrapper functions and PHMM setup
##################
nt_start = 250
nt_end = 250+395+3
aa_start = ceiling(nt_start/3) 
aa_end = ceiling(nt_end/3)

phmm_nt_250_start = coil::subsetPHMM(coil::nt_coi_PHMM, nt_start, nt_end)
phmm_aa_250_start = coil::subsetPHMM(coil::aa_coi_PHMM, start = aa_start, end = aa_end)


x = coil::coi5p_pipe(test$seq[[2]], nt_PHMM = phmm_nt_250_start,
									aa_PHMM = phmm_aa_250_start)
x$data$ntPath
x


meta_coil_check = function(seq, trans_table = 5,
				nt_PHMM = phmm_nt_250_start, aa_PHMM = phmm_aa_250_start){

	x = coil::coi5p_pipe(seq, trans_table = trans_table, 
						nt_PHMM = nt_PHMM, aa_PHMM = aa_PHMM)

	out = coil::flatten_coi5p(list(x))

	return(out)
}


get_centroid_coil_data = function(seq_df){

	centroid = seq_df[seq_df$centroid == TRUE,]

	coil_out = meta_coil_check(centroid$seq)

	return(cbind(centroid, coil_out))

}

get_mbrave_centroid_coil_data = function(seq_df){

	centroid = seq_df[seq_df$original_centroid == TRUE,]

	coil_out = meta_coil_check(centroid$seq)

	return(cbind(centroid, coil_out))

}


#get the consensus sequence using debar's drop down technique,
# returns the flattend coil data bound to the original centroid row
# so 'raw' is the consensus sequence generated by debar
get_denoised_consensus_data = function(seq_df, max_len = 396){
	print(seq_df$otu[[1]])
	centroid = seq_df[seq_df$centroid == TRUE,]

	seq_df = seq_df[nchar(seq_df$denoised_seqs) <= max_len,]
	seq_df = seq_df[!is.null(seq_df$denoised_seqs),]	
	seq_df = seq_df[!is.na(seq_df$denoised_seqs),]

	if(length(seq_df$denoised_seqs) == 0){
		fake_row = list("name"=NA, "raw"=NA, "framed"=NA, "was_trimmed"=NA,
			"align_report"=NA,"aaSeq"=NA,"aaScore"=NA, "indel_likely"=NA,"stop_codons"=NA)
		con_out = data.frame(fake_row)

	}else{
		con_seq = debar::consensus_sequence(seq_df$denoised_seqs)
		con_out = meta_coil_check(con_seq)
	}

	return(cbind(centroid, con_out))
}


has_n = function(seq){
	return(grepl('n', seq, ignore.case = TRUE))
}


count_n = function(seq){
	return(lengths(regmatches(seq, gregexpr("n", seq))))
}


######
# BASELINES
######
# centroid analysis - named OTUs

named_centroid_coil_outputs = lapply(split_named_otus, get_centroid_coil_data)
unnamed_centroid_coil_outputs = lapply(split_unnamed_otus, get_centroid_coil_data)
mBRAVE_centroid_coil_outputs = lapply(split_unnamed_otus, get_mbrave_centroid_coil_data)

named_centroid_coil_df = do.call("rbind", named_centroid_coil_outputs)
unnamed_centroid_coil_df = do.call("rbind", unnamed_centroid_coil_outputs)
mBRAVE_centroid_coil_df = do.call("rbind", mBRAVE_centroid_coil_outputs)

write_tsv(named_centroid_coil_df, 'data/named_otus_coil_baseline.tsv')
write_tsv(unnamed_centroid_coil_df, 'data/unnamed_otus_coil_baseline.tsv')
write_tsv(mBRAVE_centroid_coil_df, 'data/mBRAVE_otus_coil_baseline.tsv')

#NAMED
sum(named_centroid_coil_df$stop_codons) #125 centroids with stop codon!
sum(named_centroid_coil_df$stop_codons)/length(named_centroid_coil_df$stop_codons)
#31.4 % of total

#UNNAMED
sum(unnamed_centroid_coil_df$stop_codons) #677 centroids with stop codon!
sum(unnamed_centroid_coil_df$stop_codons)/length(unnamed_centroid_coil_df$stop_codons)
#53% with stop codon

#MBRAVE CENTROIDS
sum(mBRAVE_centroid_coil_df$stop_codons) #681 centroids with stop codon!
sum(mBRAVE_centroid_coil_df$stop_codons)/length(mBRAVE_centroid_coil_df$stop_codons)
#54% of mBrave centroids had stop codon.

mBRAVE_centroid_coil_df = as_tibble(mBRAVE_centroid_coil_df)


##########
# consensus with denoised reads
##########
named_debar_consensus = lapply(split_named_otus, get_denoised_consensus_data)
unnamed_debar_consensus = lapply(split_unnamed_otus, get_denoised_consensus_data)

named_debar_consensus_df = do.call("rbind", named_debar_consensus)
unnamed_debar_consensus_df = do.call("rbind", unnamed_debar_consensus)


named_debar_consensus_df$read_count = unlist(lapply(split_named_otus, function(x){
	nrow(x)
	}))

named_debar_consensus_df$contains_n = unlist(lapply(named_debar_consensus_df$raw, function(x){
	has_n(x)
	}))

named_debar_consensus_df$n_count = unlist(lapply(named_debar_consensus_df$raw, function(x){
	count_n(x)
	}))


unnamed_debar_consensus_df$read_count = unlist(lapply(split_unnamed_otus, function(x){
	nrow(x)
	}))

unnamed_debar_consensus_df$contains_n = unlist(lapply(unnamed_debar_consensus_df$raw, function(x){
	has_n(x)
	}))

unnamed_debar_consensus_df$n_count = unlist(lapply(unnamed_debar_consensus_df$raw, function(x){
	count_n(x)
	}))


#check individual ones
#seq_df = split_unnamed_otus[["OTU1005"]]

#NAMED - numbers are exact same>
#398 total named otu consensus sequences
sum(named_debar_consensus_df$stop_codons) #7 centroids with stop codon!
sum(named_debar_consensus_df$stop_codons)/length(named_debar_consensus_df$stop_codons)
#1.8 % of total

#^GET NUMBER OF NAs - none for the named sequences
sum(is.na(named_debar_consensus_df$raw))

# GET THE AMOUNT OF MISSING INFORMATION PER BARCODE (Ns in final)
sum(named_debar_consensus_df$contains_n)
hist(named_debar_consensus_df$n_count)

mean(named_debar_consensus_df$n_count)
mean(named_debar_consensus_df$n_count[named_debar_consensus_df$n_count!=0])

median(named_debar_consensus_df$n_count)
median(named_debar_consensus_df$n_count[named_debar_consensus_df$n_count!=0])

sum(named_debar_consensus_df$n_count>15)


#UNNAMED - numbers are exact same>
#1255 total sequences
sum(unnamed_debar_consensus_df$stop_codons, na.rm=TRUE) #128 consensus with stop codon!
sum(unnamed_debar_consensus_df$stop_codons, na.rm=TRUE)/length(unnamed_debar_consensus_df$stop_codons)
#10.2 % of total

#^GET NUMBER OF NAs #69 NA
sum(is.na(unnamed_debar_consensus_df$raw))
sum(is.na(unnamed_debar_consensus_df$raw))/length(unnamed_debar_consensus_df$stop_codons)
#5.5% failed the debar check or basic length QC

# GET THE AMOUNT OF MISSING INFORMATION PER BARCODE (Ns in final)
sum(unnamed_debar_consensus_df$contains_n)
hist(unnamed_debar_consensus_df$n_count)

mean(unnamed_debar_consensus_df$n_count)
mean(unnamed_debar_consensus_df$n_count[unnamed_debar_consensus_df$n_count!=0])

median(unnamed_debar_consensus_df$n_count)
median(unnamed_debar_consensus_df$n_count[unnamed_debar_consensus_df$n_count!=0])

sum(unnamed_debar_consensus_df$n_count>15)


#####################################
# write dfs to file.

write_tsv(named_debar_consensus_df, 'data/named_otus_debar_consensus_seqs.tsv')
write_tsv(unnamed_debar_consensus_df, 'data/unnamed_otus_debar_consensus_seqs.tsv')


#######################
# Sequence-by-sequence analysis
#######################
trans_table = 5
nt_PHMM = phmm_nt_250_start
aa_PHMM = phmm_aa_250_start

seq = unnamed_otu_data$seq[[2193]]

x = coil::coi5p_pipe(seq, trans_table = trans_table, 
					nt_PHMM = nt_PHMM, aa_PHMM = aa_PHMM)

out = coil::flatten_coi5p(list(x))


#slow version, b/c failure with mclapply
mBRAVE_original_coil_check = lapply(1:nrow(unnamed_otu_data), function(x){
	print(unnamed_otu_data$label[[x]])
	meta_coil_check(unnamed_otu_data$seq[[x]])
})


df_mBRAVE_original_coil_check = do.call("rbind", mBRAVE_original_coil_check)

mBRAVE_original_coil_output = cbind(unnamed_otu_data, df_mBRAVE_original_coil_check)

write_tsv(mBRAVE_original_coil_output, 'data/original_OTU_seqbyseq_coil.tsv')


mBRAVE_denoised_coil_check = mclapply(unnamed_otu_data$denoised_seqs,
							meta_coil_check, mc.cores = detectCores())


#NAs cause coil failure, remove them before the denoised test.
temp = unnamed_otu_data[unnamed_otu_data$label == "423YV:02423:04030",]

unnamed_otu_data = unnamed_otu_data[!is.null(unnamed_otu_data$denoised_seqs),]	
unnamed_otu_data = unnamed_otu_data[!is.na(unnamed_otu_data$denoised_seqs),]


#slow version to avoid issue with the mclapply call
mBRAVE_denoised_coil_check = lapply(1:nrow(unnamed_otu_data), function(x){
	print(unnamed_otu_data$label[[x]])
	meta_coil_check(unnamed_otu_data$denoised_seqs[[x]])
})


df_mBRAVE_denoised_coil_check = do.call("rbind", mBRAVE_denoised_coil_check)

mBRAVE_denoised_coil_output = cbind(unnamed_otu_data, df_mBRAVE_denoised_coil_check)

write_tsv(mBRAVE_denoised_coil_output, 'data/denoised_OTU_seqbyseq_coil.tsv')
```
