## Supplementary material for "debar, a sequence-by-sequence denoiser for COI-5P DNA barcode data": debar_supplementary: denoising_reads_keep_right.r

```
print("loading libraries")
library(coil)
library(debar)
library(parallel)
library(tidyverse)

print("loading data")
data = read_tsv('data/run_18333_derep_data.tsv')
#for small test
#data = head(data)

print("subsetting the PHMM")
######
#subset the PHMM for the metabarcode region
###### 
nt_start = 250
nt_end = 250+395+3
phmm_nt_250_start = coil::subsetPHMM(debar::nt_coi_PHMM, nt_start, nt_end)

print("detecting cores")
ncores = parallel::detectCores()

print("denoise the sequences")

#data = data[1:100,]

#denoise the sequences, add to the df
denoised_seqs = debar::denoise_list(data$seq, 
						cores = ncores,
						nt_PHMM = phmm_nt_250_start,
						keep_flanks = 'right',
						dir_check = FALSE)

#x = data$seq[[99]]
#dat = DNAseq(x)
#dat = frame(dat, nt_PHMM = phmm_nt_250_start, dir_check = FALSE)
#y = denoise(x,
#	nt_PHMM = phmm_nt_250_start,
#						keep_flanks = 'right')
#dat$data$path

data$denoised_seqs = unlist(lapply(denoised_seqs, function(x){
			if(is.null(x)){
				return(NA)
			}else{
				return(x)
			}}))

print("writing to file")
write_tsv(data, 'data/denoised_run_18333_derep_data_keepright.tsv')
```
