## Supplementary material for "debar, a sequence-by-sequence denoiser for COI-5P DNA barcode data": debar_supplementary: S5-debar-vignette.pdf

### debar - DEnoising COI-5P BARcode data

Cameron M. Nugent

```r
library(debar)
```

#### Introduction

`debar` is an R package designed for denoising high throughput sequencing data for the animal DNA barcode marker cytochrome c oxidase I (COI-5P, or the five prime portion of COI). The package is designed to detect and correct insertion and deletion errors within sequencer outputs. This is accomplished through comparison of input sequences against a profile hidden Markov model using the Viterbi algorithm (`debar` depends on functions from the R package aphid for running the Viterbi algorithm). Inserted base pairs are removed and deleted base pairs are accounted for through the introduction of a placeholder character. Since the PHMM is a probabilistic representation of the COI barcode, corrections are not always perfect. For this reason `debar` censors base pairs adjacent to reported indel sites, turning them into placeholder characters (default is 7bp in either direction, this feature can be disabled). Testing has shown that this censorship results in the correct sequence length being restored, and erroneous base pairs being masked the vast majority of the time (>95%).

##### Implementation

The package is designed to detect and correct insertion and deletion errors within sequencer outputs. This is accomplished through comparison of input sequences against a profile hidden Markov model using the Viterbi algorithm. The Viterbi path produced by this comparison is used to identify where insertion or deletion errors exist. Here is a small example of how the package corrects indels in this manner:

```
#Correcting insertions
#the 'G' in this sequence is an insertion
"ATCATGATC"
#the sequence is compared to the PHMM, producing a Viterbi path
#the 'G' is assigned a hidden state of 'insert', indicated by a 2 in the path
"111112111"
#debar uses this information to alter the sequence, removing the G
"ATGATATG"
```

For deletion errors the inverse action is performed and a placeholder character is inserted.

##### Dependencies

`debar` is dependent on the `aphid` package for comparison of sequences against a COI-5P nucleotide PHMM. The package `ape` is a requirement because sequences are internally converted to `ape` "DNAbin" objects. The package `sequinr` is utilized in obtaining the reverse compliment of input sequences when needed. The `parallel` package is also required for parallelization of the denoising process.

#### Using the package

##### Denoising a list of sequences

When working with sequence data in R, `debar` can be applied to a set of reads (either those linked to a given sample or those from a given set of clustered sequences). Note that for best performance reads should already be demultiplexed, dereplicated, and filtered based on quality. A list of sequences can be denoised with the `denoise_list` function.

```
ex_out = denoise_list(ex_nt_list)
```

`denoise_list` operates on a sequence-by-sequence basis and is therefore parallelized to improve performance. The `cores` argument allows a user to specify the number of cores across which the processes should be parallelized.

```
ex_out = denoise_list(ex_nt_list, cores = 4)
```

Passing the option `keep_flanks=FALSE` to the `denoise_list` function will put all of the outputs into a common reading frame. This allows for the output lists to be combined to produce a consensus sequence.

```
ex_out = denoise_list(ex_nt_list, keep_flanks=FALSE)
```

##### Obtaining a consensus sequence

`debar` can provide a more accurate consensus sequence for a sample by removing errors in component reads and ensuring that all data conform to the known amplicon structure. The denoising process yields a series of indel corrected sequence reads, which can be combined to obtain a biologically informed consensus if the reads are derived from a single sample or from a sufficiently similar set of metabarcode outputs. The censorship process effectively removes indel errors the majority of the time and also restores the proper length of the sequence. This comes at the cost of missing information in individual sequences. The combination of denoised sequences from a given sample or amplicon variant allows the full barcode to be obtained in most instances using the `consensus_sequence` function.

```
barcode_out = consensus_sequence(ex_out)
```

```
# truncated example of debar outputs for a given sample
# indels corrected and area masked with 'N' placeholders
CTCTACTTGNNNNNNNNNNNNNNNAGCAGGAATAGTTGGAATAGCTTTAAGTTTACTAATTCGCGCTGAACTAGGT
CTNNNNNNNNNNNNNNNNTGCATGAGCAGGAATAGTTGGAATAGCTTTAAGTTTACTAATTCGCGCTGAACTAGGT
CTCTACTTGATTTTTGGTGCATGAGCAGGAATAGTTGGAATAGCTTTAAGTTTNNNNNNNNNNNNNNAACTAGGT

# following assignment to samples (barcoding) or OTU clustering (metabarcoding)
# a denoised barcode can be obtained from the consensus of the denoised reads

CTCTACTTGATTTTTGGTGCATGAGCAGGAATAGTTGGAATAGCTTTAAGTTTACTAATTCGCGCTGAACTAGGT
```

Note: If you are interested in identifying which consensus sequences do not conform to the COI amplicon structure, as opposed to denoising the sequences, then consider employing `debar`s sister package: `coil`

##### File-to-file denoising

The program contains a wrapper function `denoise_file` that allows for a user to run the entire denoiser pipeline in a single step (there are quantitatively informed defaults for executing the full pipeline, but it is highly recommended that a user acclimate themselves to the `denoise` pipeline options to ensure that the parameters make sense for their given data set and analysis goals).

All that is needed is for the input and output files to be specified. The program by default will read in the input file, run each sequence through the denoising algorithm, and then output the denoised sequences to the specified output file. The denoise pipeline contains certain rules (that a user can control - see below for details on these parameters) that lead to some sequences being rejected entirely, the `keep_rejects` option allows a user to output these rejected sequences to another file for further inspection. The program also has an option for generating a simple log file, and allows for multicore execution of the denoising pipeline by simply specify the number of cores the process should utilize. Additionally you can type `?denoise` to see a list of all available denoise parameters that can be passed to the `denoise_file` wrapper function as well (parameters are explored in more detail below).

##### Data structure

`debar` can operate on inputs from fasta and fastq files, or their gzipped equivalents. For demonstration of the workflow, `debar` contains three example files.

```
#The following example file is used as an input in the vignette
fastq_example_file = system.file('extdata/coi_sequel_data_subset.fastq.gz',
                                 package = 'debar')

#other example data also available in debar:
#fastq_example_file = system.file('extdata/coi_sequel_data_subset.fastq',
#                                 package = 'debar')
#fasta_example_file = system.file('extdata/coi_sequel_data_subset.fasta',
#                                 package = 'debar')
```

##### File-to-file example

```
# NOTE - this block of code is not run so as to avoid the generation of output files!
#
# Our input file with noisy sequence data is fastq_example_file.
# Output file is "example_output.fastq"
# If you do wish to run these examples, then please double check your working directory!
denoise_file(fastq_example_file, filename = "example_output.fastq")

denoise_file(fastq_example_file, filename = "multicore-example_output.fastq",
                          multicore = 8, log_file = TRUE, keep_rejects = TRUE)
```

##### Parameter combinations - speed and accuracy trade offs

File-to-file denoising can also be parallelized across multiple CPU cores. The denoising of each sequences in the input file is conducted separately, so using multiple cores will decrease the time needed to complete denoising roughly by a factor of the number of cores used. If you are denoising complete sequencer outputs, it is highly recommended that you do so with as many cores as possible. For example, denoising of 160,000 sequence reads on a 64-core server (all default parameters) takes approximately 1hr and 42 minutes, while on a laptop with only 8 cores would take almost 14 hours!

```
#debar works best when the tasks are highly parallelized
denoise_file(fastq_example_file, filename = "multicore-example_output.fastq",
             multicore = 8, log_file = TRUE, keep_rejects = TRUE)
#set the multicore parameter to the number of CPU cores available
```

Certain parameter selections can further increase the speed with which `debar` can process data, but come with certain trade offs (that may or may not be worth consideration in the processing of your own data). The most drastic speed improvement is provided by disabling the direction check (dir_check). By default both the forward and reverse compliments of a read are compared to the PHMM, if your data consists only of forward reads, then disabling this option will result in a 30-50% reduction in processing. Other speed/accuracy trade offs are available and discussed within the 'Recommended parameter combinations' section of the package's vignette.

##### Denoising metabarcode data

DNA metabarcoding analyses often target shorter, standardized sections of the full COI barcode region for amplification and sequencing. If the targeted section is known, the performance of debar can be optimized by denoising the sequence using a customized subset of the complete PHMM, corresponding to given region. This will reduce the probability of spurious profile matches, and increase the accuracy of the indel corrections performed.

debar's sister package, coil contains a function `subsetPHMM`, which can be used to extract a section of the complete 657bp PHMM for the COI region. Below this process, and the application of the custom model in the denoising of a barcode sequence fragment are demonstrated.

```
#install.packages('coil')

# These variables define the region of the Folmer region we are targeting.
nt_start = 295
nt_end = 636

# Take the complete PHMM, and create a new PHMM that represents
#a specific subsection of the COI Folmer region.
subsection_phmm = coil::subsetPHMM(debar::nt_coi_PHMM, start = nt_start, end = nt_end)

# Here is an example sequence with a T added at position 81.
ex_error = paste0(c("tccgcaggagtagaagctggagcaggaaccggatgaactg",
"tatatcctcctttagcaggtaatttagcacatgctggccacctctgttgatttagccatctttt",
"cccttcatttggccggtatctcatcaattttagcctctattaattttattacaactattatta",
"atataaaaccccaactatttctcaatatcaaacaccattatttgtttgatctattcttatcac",
"cactgttcttctactccttgctctccctgttcttgcagccggaattacaatattattaacagacc",
"gcaacctcaacactacattctttgaccccgcaggggggaggggacc"), collapse = "")

# Run the denoise function, with the constrained PHMM.
subset_has_error_outpt = denoise(ex_error, nt_PHMM = subsection_phmm)

subset_has_error_outpt
```

```
## A DNAseq object.
## Raw Sequence:
## tccgcaggagtagaagctggagcag...tttgaccccgcaggggggaggggacc
## Corrections applied: 1
## Output Sequence:
## TCCGCAGGAGTAGAAGCTGGAGCAG...TTTGACCCCGCAGGGGGGAGGGGACC
```

#### Acknowledgements

Funding for the development of this software was provided by grants in Bioinformatics and Computational Biology from the Government of Canada through Genome Canada and Ontario Genomics and from the Ontario Research Fund. Funders played no role in the study design or preparation of this software.
